# Reciprocal feedback inhibition and divergent intrinsic firing properties between behaviorally defined classes of neurons in the mouse ventromedial hypothalamus

**DOI:** 10.64898/2026.07.30.741940

**Authors:** Sukrita Deb, Sofia Torchia, Emily Welponer, Luca Spagnoletti, Selin Karagülle, Sofia Aravantis, Maria Esteban Masferrer, Yerassyl Zhalgasbayev, Hiroki Asari, Cornelius T. Gross

## Abstract

The capacity to orchestrate appropriate defensive behaviors in response to threatening stimuli is essential for the survival of organisms. Flight, freezing, appeasement, and attack, for example, are common defensive responses evoked depending upon the nature of the threat, the immediate context, and past experiences. Multiple brain regions have been implicated in regulating such responses, with the medial hypothalamus playing a key role in mediating innate responses to social and predator threats. The ventrolateral subdivision of the ventromedial hypothalamus (VMHvl) has been shown to mediate both avoidance and aggression towards conspecific threats in mice, suggesting that it serves as a circuit for context-appropriate defensive responses to social threats. Previous *in vivo* calcium imaging in VMHvl identified cells whose activity encodes either approach toward or escape from a social threat (*Assessment*+ and *Flight*+ cells, respectively), but it remained unclear how the switch in neural encoding from approach to avoidance occurs. Here, we use *in vivo* single-unit electrophysiology recordings coupled to channelrhodopsin-assisted circuit mapping (optrodes) to explore the functional and structural connectivity basis of the approach-avoidance switch. We confirm the presence of *Assessment*+ and *Flight*+ neurons in VMHvl and demonstrate that they are interconnected through feedback excitation and reciprocal feedback inhibition. Moreover, we discover that *Assessment*+ and *Flight*+ neurons exhibit a pronounced difference in their intrinsic firing properties. We hypothesize that this asymmetry in intrinsic responsivity coupled to reciprocal feedback inhibition underlies the nonlinear firing changes at the approach-to-avoidance transition and plays a role in triggering escape behavior.

## Introduction

Defensive behaviors are a crucial part of the repertoire that allow organisms to adapt to threats, with several brain regions implicated in their expression. Multiple interconnected nuclei in the mammalian diencephalon have been proposed to form the medial hypothalamic defensive system (Canteras, 2002) and both anatomical and functional studies have given support to the existence of two parallel pathways within this system dedicated to the control of defensive behaviors to predator versus social threats (Gross and Canteras, 2012; Silva et al., 2013). Within the medial hypothalamic defensive system, the ventromedial hypothalamus (VMH) is strategically positioned because it receives converging inputs carrying multi-modal sensory information about threats from the amygdalar complex and contextual information from the hippocampus via the lateral septum, while promoting defensive behavior and autonomic changes via outputs to the periaqueductal grey (Canteras et al., 1994; Wang et al., 2015; Lo et al., 2019; Montardy et al., 2020). Its dorsomedial subdivision (VMHdm) forms part of the predator defense pathway, while the ventrolateral subdivision (VMHvl) forms part of the social defense pathway (Gross and Canteras, 2012; Silva et al., 2013). Interestingly, neuronal activity in VMHvl is required for the production of both escape responses to a social threat and attacks toward such threats, suggesting that the medial hypothalamus plays a general role in supporting context-appropriate social threat responding regardless of whether that involves attacking an intruder or escaping from a bully, for example (Lin et al., 2011; Sakurai et al., 2016; Hashikawa et al., 2017; Wang et al., 2019; Falkner et al., 2020; Krzywkowski et al., 2020). Likewise, artificial activation of VMHvl excitatory core neurons is able to elicit both social avoidance and attack, depending on context and experience (Lin et al., 2011; Lee et al., 2014; Sakurai et al., 2016; Wang et al., 2019; Krzywkowski et al., 2020; Sureka et al., 2025) with some evidence that these responses may segregate across the anterior-posterior axis of the nucleus (Wang et al., 2019; but see Lee et al., 2014; Krzywkowski et al., 2020).

However, the local circuit mechanisms by which VMHvl might trigger the decision to attack or escape when encountering a social threat remain unclear. One clue comes from work that has shown that repeated experience of attacking another mouse or being attacked by a bully can reprogram synaptic connectivity in VMH in a manner that biases future social encounters in favor of social aggression or avoidance, respectively (Lee et al., 2014; Krzywkowski et al., 2020; Stagkourakis et al., 2020). However, we lack an understanding of how this plasticity impacts the processing of threat information in VMHvl because we lack a model of the synaptic connectivity and input-output transformations that underlie the contribution of VMHvl to triggering escape or attack behaviors.

Single-unit neural recordings in VMH during defensive behaviors have identified distinct classes of neurons with unique behavioral response properties and have permitted the construction of the first local circuit models for VMH function (MacGregor and Leng, 2019; Kennedy et al., 2020; Rahy et al., 2022; Alfieri et al., 2022; Nair et al., 2023). In a pioneering study, VMH single-units were recorded for the first time in male mice while they approached and attacked an intruder (Lin et al., 2011). Two classes of neurons were identified which increased or decreased, respectively, their firing as the animal approached the intruder (*Attack*+ and *Attack*- neurons). Notably, the probability of attack could be predicted from the magnitude of firing of *Attack*+ neurons during the approach phase, and *Attack*+ neuron activity persisted as the attack progressed (Lin et al., 2011; Falkner et al., 2014) suggesting that VMH activity dictates the probability of attack, but does not encode the decision to attack, which presumably occurs in downstream brain structures.

On the other hand, when VMH single-units were recorded in mice that approached and escaped from a threat, two classes of neurons were identified (Esteban Masferrer et al., 2020; Krzywkowski et al., 2020). The first class increased their activity during the approach toward the threat, much like *Attack*+ neurons, and were called *Assessment*+ neurons. However, unlike *Attack*+ neurons, the firing of *Assessment*+ neurons abruptly ceased upon onset of escape, suggesting that they might encode the decision to escape. The second class showed little modulation during approach, but abruptly increased their firing at the onset of escape, and were called *Flight*+ neurons. Collectively, this suggests that the reciprocal firing changes in *Assessment*+ and *Flight*+ neurons encode the escape decision point, and that, while VMHvl firing activity may be restricted to influencing the probability of attack during aggression, it has a more direct role in triggering escape during defense. A causal role for VMHvl activity in triggering escape is supported by studies in which optogenetic activation of VMHvl evokes flight or avoidance (Sakurai et al., 2016; Krzywkowski et al., 2020).

However, understanding whether VMHvl is involved in setting the escape decision threshold would require understanding how local circuitry supports the abrupt, nonlinear changes in activity of *Assessment*+ and *Flight*+ neurons. An exploration of local connectivity and synaptic plasticity parameters in a computational model of VMH found that asymmetric reciprocal feedback inhibition between *Assessment*+ and *Flight*+ neurons is sufficient to recreate the simultaneous decrease and increase in firing of these neuron classes seen at escape onset (Rahy et al., 2022). Intriguingly, while core VMH neurons are excitatory, surrounding VMH shell neurons are inhibitory (Yamamoto et al., 2018; Kim et al., 2019), raising the possibility of local reciprocal feedback inhibition between core VMH neurons via GABAergic shell neurons.

In this study, we tested the hypothesis that local circuitry features in VMHvl may support the decision to escape by investigating the activity and connectivity of VMHvl neurons using *in vivo* single-unit electrophysiology and optrode-mediated circuit mapping (Lima et al., 2009). We first developed a novel behavioral test that allowed us to record VMHvl neuron activity during unrestricted social approach and escape between male mice. Using this test, we confirmed the presence of *Assessment*+ and *Flight*+ neurons in VMHvl and found that *Flight*+ neuron activity predicted the vigour of escape behavior on a trial-by-trial basis. Using optrode neuron-tagging we recorded from GABAergic VMHvl shell neurons and found that they too harbor *Assessment*+ and *Flight*+ neurons, as predicted by the reciprocal feedback inhibition model. Finally, we used *in vivo* channelrhodopsin-evoked somatic and synaptic stimulation to infer the connectivity and intrinsic firing properties of *Assessment*+ and *Flight*+ neurons. Our findings demonstrate the presence of reciprocal feedback inhibition between *Assessment*+ and *Flight*+ neurons in VMHvl and document pronounced differences in their intrinsic firing properties, offering support for a role of VMH in local escape decision calculation. Together, these findings provide a local circuitry basis for the activity patterns observed in VMH and make predictions about how this structure can control social threat response thresholds.

## Results

### Establishment of a social pursuit testing apparatus

Current behavioral tests for social avoidance typically use small arenas or the home cage which do not allow for sustained flight and unfettered pursuit between animals (Lin et al., 2011; Silva et al., 2013). To allow for *in vivo* neural recordings during more naturalistic and sustained social escape behaviors, we designed a novel circular arena that allowed unlimited social pursuit between pairs of animals (**Figure 1A**, **Figure S1A**; Griebel et al., 1996). The behavioral protocol primarily consisted of two phases of interest when social interactions were scored (**Figure 1B**). In the first Restricted Interaction (RI) phase, an aggressive male CD1 outbred mouse was confined in a wiremesh cage in a small section of the circular corridor. An experimental male C57BL/6J mouse was then placed into the apparatus so that it could repeatedly approach and investigate the threat from either side. Frequently, the experimental mouse turned and fled from the aggressor after a period of investigation (**Figure 1C**, **Figure S1B**), allowing for the quantification of repeated approach-escape trials (**Figure 1D**).

**Figure 1.**
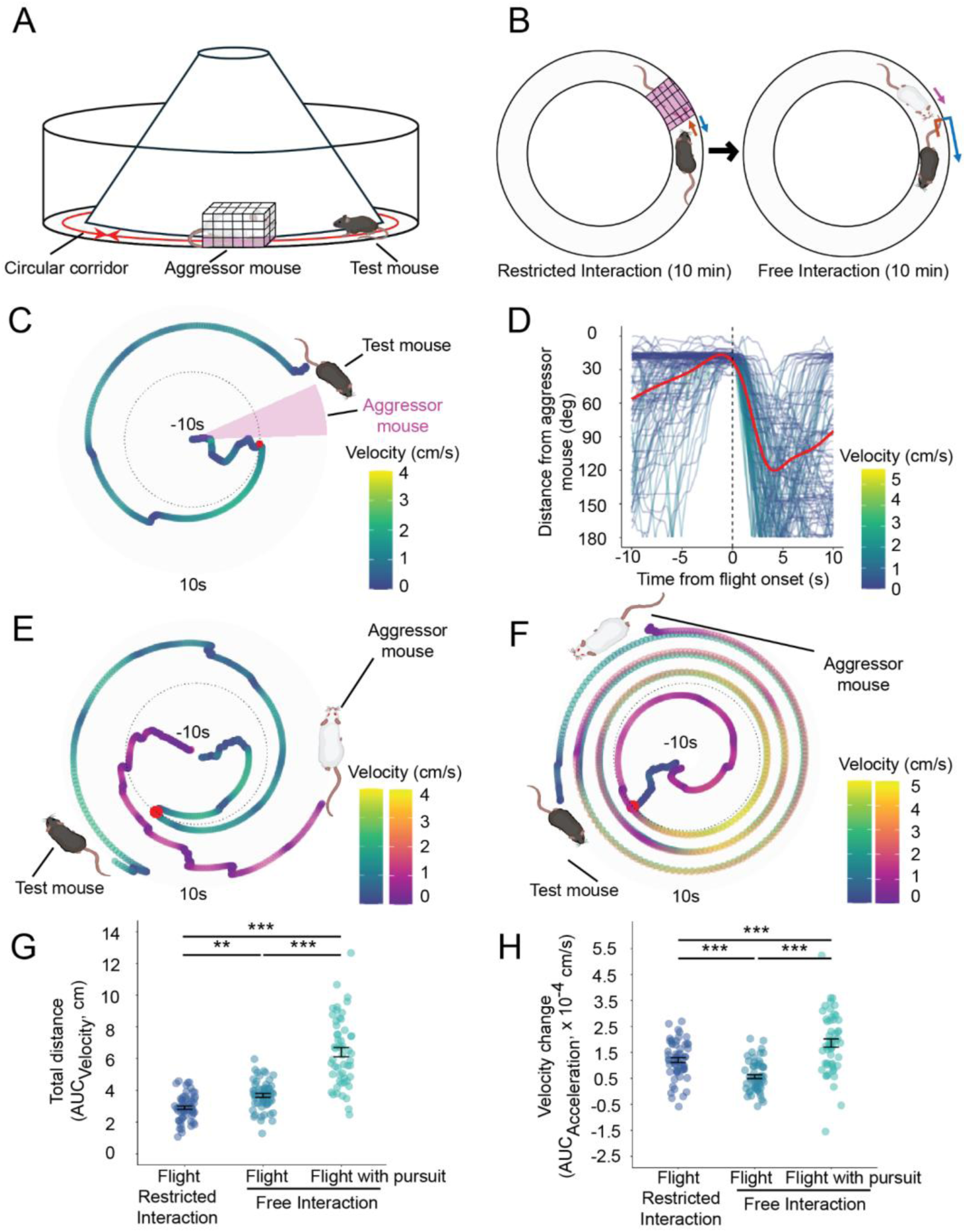
Social pursuit test. ***A***, Cartoon of the circular arena seen from the side. ***B***, The phases of the behavioral protocol – Restricted interaction (RI) and Free interaction (FI). ***C***, Representative example of an assessment-escape trial in the RI phase carried out towards the caged aggressor (pink sector). The azimuth angle of the experimental mouse coloured according to its velocity in the period from -10 (center of the circle) to +10 s (circumference of the circle) centered on flight onset (red dot) at time 0 s (dotted circle). ***D***, Multiple trials of assessment-escape depicting the change in distance (in degrees) between the experimental mouse and the aggressor in the -10 to +10 s period from flight onset coloured according to movement velocity with the predicted trend in red (N = 103 trials). ***E***, Representative example of an assessment-escape trial in the FI phase carried out towards the freely moving aggressor. ***F***, Representative example of a flight with pursuit trial in the FI phase. ***G***, Quantification of the change in position of the animal in a trial for flights in RI (N = 59), flights in FI (N = 58) and flights with pursuit in FI (N = 54). Each point in a group is an average of the trial measures in a session (ANOVA F = 96.506, *p < 0.001*, Tukey’s flight_RI_ vs flight_FI_ *p = 0.009,* flight_RI_ vs flight with pursuit *p < 0.001*; flight_FI_ vs flight with pursuit *p < 0.001*; error bars represent mean ± SEM). ***H***, Quantification of the change in velocity of the animal in a trial for flights in RI (N = 59), flights in FI (N = 58) and flights with pursuit in FI (N = 54). Each point in a group is an average of the trial measures in a session (ANOVA F = 32.589, *p < 0.001*, Tukey’s flight_RI_ vs flight_FI_ *p < 0.001,* flight_RI_ vs flight with pursuit *p < 0.001*; flight_FI_ vs flight with pursuit *p < 0.001*; error bars represent mean ± SEM; * *p < 0.05, ** p < 0.01, *** p < 0.001*).

The Restricted Interaction phase was followed by a Free Interaction (FI) phase where the wire mesh cage was removed and both mice were able to explore the apparatus freely. Under these conditions the experimental mouse was frequently pursued by the aggressor and exhibited more prolonged flights (**Figure 1E, F**; **Figure S1C, D**). A quantification across multiple experimental sessions and animals confirmed a significantly greater distance moved initially after behavior onset during Free vs. Restricted Interaction, and during flight with vs. without pursuit (**Figure 1G**). The early velocity change during the flight trials, on the other hand, was significantly lower in Free vs. Restricted Interaction phases, but increased with pursuit (**Figure 1H**). The reduction in the change in flight velocity during the Free Interaction phase may reflect less perseverative escapes, with greater variation in acceleration under these conditions. Moreover, the time spent in close investigation was significantly decreased in the Free Interaction phase (**Figure S1G**). As expected, the duration of flights was significantly longer during flight with pursuit than under other conditions (**Figure S1I**) and the mean velocity of flights was significantly greater in the free vs. restricted interaction phases and between with vs. without pursuit (**Figure S1J**) suggesting a greater perception of threat under free interaction conditions. Finally, during the Free Interaction phase the aggressor occasionally succeeded in attacking the experimental mouse and the experimental mouse frequently displayed defensive upright postures aimed at suppressing such aggression. As a result, experimental mice spent significantly less time near the wire mesh cage during a subsequent restricted social exposure that occurred after the free interaction, than in the RI phase prior (**Figure S1E-F, H**).

### Encoding of approach-escape decision in VMHvl

Using *in vivo* single-unit electrophysiology with either traditional wire-bundle or silicon probes, a total of 675 neurons were recorded in VMHvl across 17 animals and 89 experimental sessions, of which 28% (187/675) showed significant modulation of their firing rates at flight onset. Behavior-responsive neurons were clustered in an unbiased manner based on their activity centered around flight onset (**Figure 2A, Figure S2A-B, D-E**) to obtain two major classes that showed *Assessment*+ (10%, 69/675) and *Flight*+ (11%, 74/675) profiles (Esteban Masferrer et al., 2020; Krzywkowski et al., 2020) and one unassigned cluster (7%, 44/675) with intermediate features to the two other classes (**Figure 2B, Figure S2C**). As expected, *Assessment*+ neurons showed a gradual increase in firing in the assessment period leading up to escape followed by a sharp decrease in firing at the onset of escape, a pattern that showed a general inverse relation to velocity (**Figure 2C, E**). Notably, the peak firing activity occurred shortly after the onset of escape, unlike what was reported previously (**Figure 2C, Figure S2F**; Esteban Masferrer et al., 2020). On the other hand, *Flight*+ neurons showed little modulation during the assessment period (**Figure S2G**), followed by a sharp increase in firing at escape onset, generally following a positive relation to velocity (**Figure 2D, F**). These *in vivo* electrophysiological observations confirm earlier calcium imaging data describing *Assessment*+ and *Flight*+ cells in VMHvl during social approach and escape (Krzywkowski et al., 2020).

**Figure 2.**
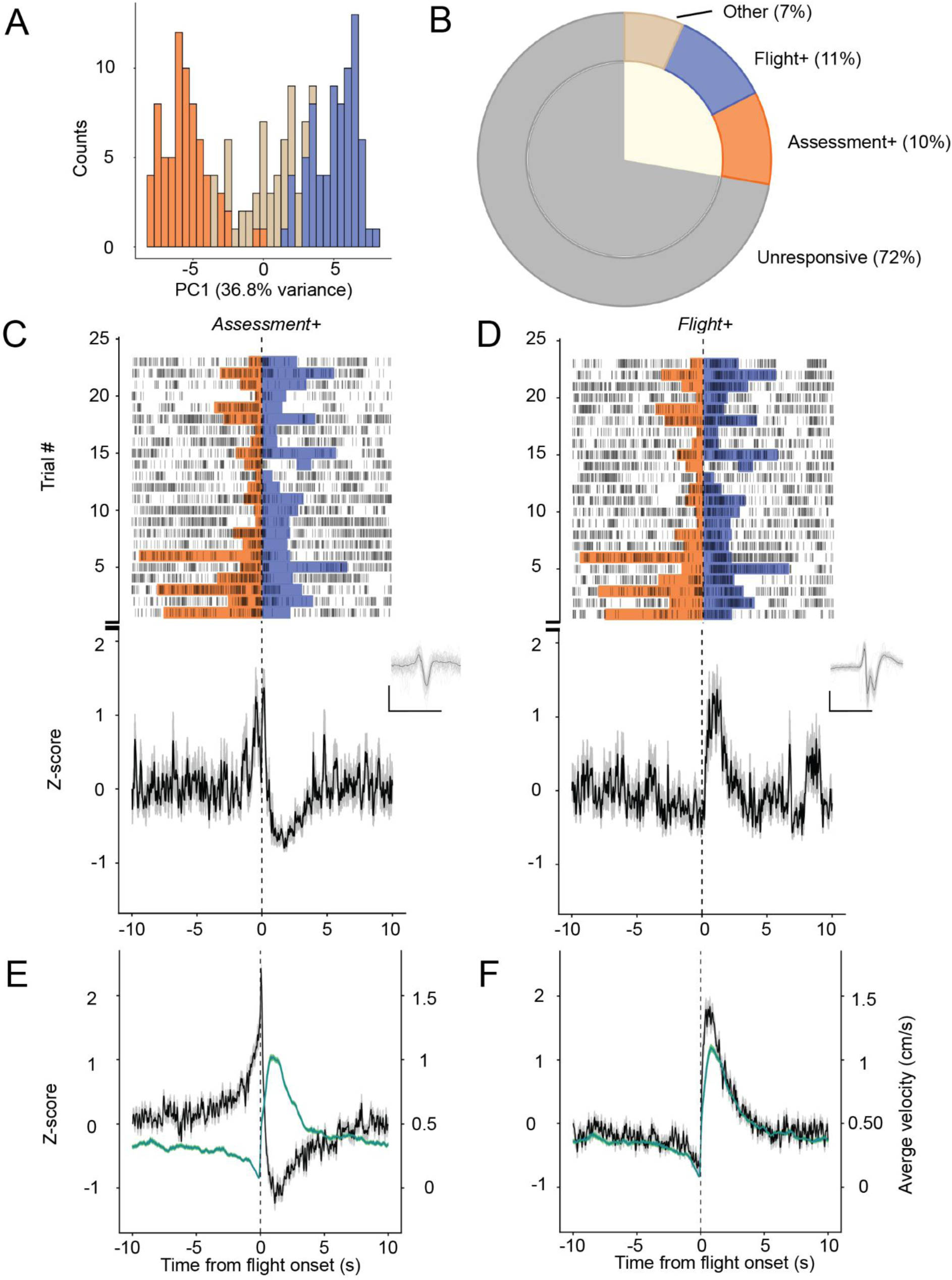
Neural activity in VMHvl during social approach-escape. ***A***, Histogram of neurons distributed according to PC1 score coloured by cluster identity and obtained by OPTICS clustering. ***B***, Population distribution of neurons found in the VMHvl (responsive neurons: mice N = 15, sessions N = 61, neurons N = 187). ***C***, Representative example of an *Assessment+* neuron. (**top**) raster plot showing firing of the neuron during transition from investigation (orange) to flight (blue), (**bottom**) average Z-scored PSTH of unit response with the grey shading indicating ±SEM, (**inset**) average and subset of traces of the waveform of the neuron (N = 50 traces, scale bar: x-axis = 1 ms, y-axis = 20 mV). ***D***, Representative example of a *Flight+* neuron. (**top**) raster plot showing firing of the unit during approach-escape transition, (**bottom**) average Z-scored PSTH of neuron response with the grey shading indicating ±SEM, (**inset**) average and subset of traces of the waveform of the neuron (N = 50 traces, scale bar: x-axis = 1 ms, y-axis = 50 mV). ***E***, Normalized average population response of *Assessment+* neurons, superposed with the average velocity of the test mice during the approach-escape transitions (N = 69 neurons from 35 sessions). ***F***, Normalized average population response of *Flight+* neurons superposed with the average velocity of the experimental mice (in green; N = 74 neurons from 34 sessions). Shading (grey for activity response, green for velocity) represents ±SEM for the respective traces.

### Flight neurons decode escape vigor

Next, we asked whether the firing of *Assessment*+ and *Flight*+ neurons was correlated with the intensity of behavior on a trial-by-trial basis. Such a correlation could be evidence for a possible causal involvement in driving behavior (i.e. does neural activity decode escape intensity?). *Flight*+ neurons were observed whose firing rates correlated significantly with the distance moved across escape trials (**Figure 3A, B**) and whose initial firing slopes correlated significantly with acceleration across trials (**Figure 3C**). Overall, the firing of 46% (34/74) of *Flight*+ neurons was significantly positively correlated with measures of escape when assessed by linear regression (**Figure 3D**). Principal component analysis (PCA) of the outputs of the linear regression analysis between *Flight*+ neuron firing and escape behavior measures (β and R^2^ values for total distance, acceleration) was used to visualize the diversity of correlations across neurons (**Figure 3E-G**). The results identified a major component (PC1, 41% variance explained) that separated correlated vs. non-correlated neurons (**Figure 3F**), and a second component (PC2, 27% variance explained) that separated neurons correlated to distance vs. acceleration (**Figure 3G**). More than 80% (28/34) of significantly correlated *Flight*+ neurons had positive PC1 scores while 47% (16/34) had positive PC2 scores (**Figure 3E**). 18/74 (24%) of the *Flight*+ neurons consistently responded to the initial velocity change of the flight, with the majority 94% (17/18) increasing their firing quantifiably with the flight velocity increase. On the other hand, the initial activity of 11% (8/74) of *Fligh*t+ neurons could predict the distance moved during the flight trial, while a further 11% (8/74) of neurons were responsive to both measures (**Figure 3D, E**). *Assessment*+ neurons, on the other hand, showed a more limited correlation with positional measures when evaluated during investigation behavior. Only 20% (14/69) of this class showed a significant correlation with the distance moved during investigation or with either the change in velocity when the animals transitioned from approach to investigation (**Figure 3H-K**). PCA identified two major components, PC1 (29% variance explained) and PC2 (26% variance explained) (**Figure 3M, N**). However, *Assessment*+ neurons did not show an interpretable distribution when projected on PCA space, consistent with their poor behavior correlation overall (**Figure 3L**). Taken together, while about half of *Flight*+ neurons decoded the vigour of escape on a trial-by-trial basis, *Assessment*+ neurons showed only a very limited correlation with behavior across trials and thus are less likely to be involved in setting the quality of behavior.

**Figure 3.**
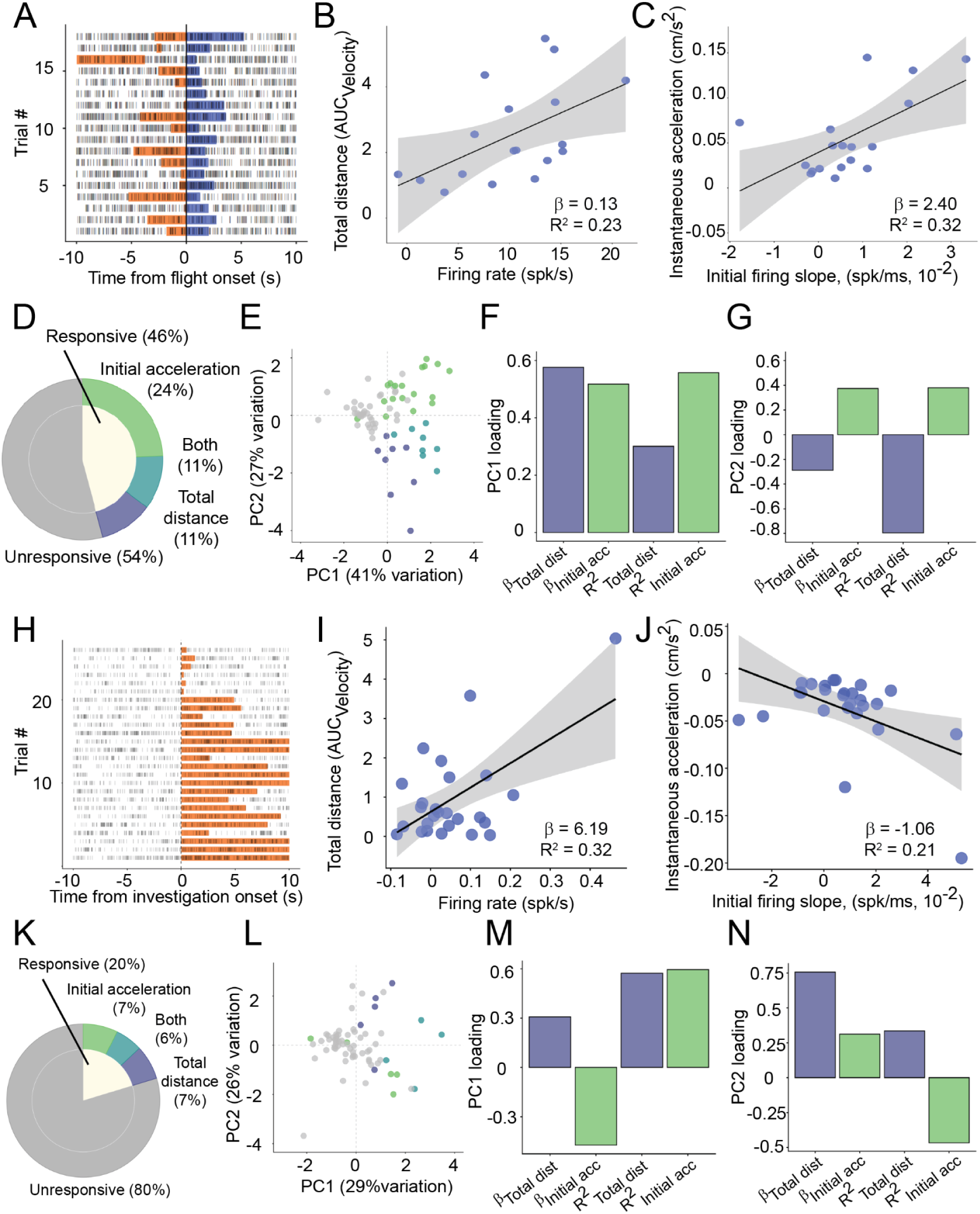
Trial-by-trial correlation between neural activity and behavior. ***A***, Raster plot showing the firing of a representative *Flight+* neuron with the onset of flight (blue) across trials. Most trials are immediately preceded by investigation (orange). ***B***, Trial-by-trial linear regression of the same unit of the total change in position of the mouse (AUC_Velocity_) during a trial against the firing in the initial 2 s period after flight onset. Regression slope β and adjusted coefficient of determination R^2^ in inset, *p_adj_ = 0.023*. Grey shading indicates 95% confidence interval. ***C***, Trial-by-trial linear regression of the change in velocity after flight onset against the firing increase in the same period (initial acceleration). Regression slope β and adjusted R^2^ in inset, *p_adj_ = 0.016*. Grey shading indicates 95% confidence interval. ***D***, Population distribution of the *Flight+* neurons responding to regression of above velocity measures against firing (N = 74 neurons). ***E***, PCA projection of the regression measures coloured according to significance of regression. ***F***, Contribution of the regression measures towards the PC1 dimension. ***G,*** Contribution of the regression measures towards the PC2 dimension. ***H,*** Raster plot showing the firing of a representative *Assessment+* neuron with the transition from approach to investigation (orange) behavior across trials. ***I,*** Trial-by-trial linear regression of the same neuron for the total change in position of the mouse (AUC_Velocity_) during a trial against the firing in the initial 2 s period after flight onset. Regression slope β and adjusted R^2^ in inset, *p_adj_ = 0.004*. Grey shading indicates 95% confidence interval. ***J,*** Trial-by-trial linear regression of the change in velocity after flight onset against the firing increase in the same period (initial acceleration). Regression slope β and adjusted R^2^ in inset, *p_adj_ = 0.011*. Grey shading indicates 95% confidence interval. ***K,*** Population distribution of the *Assessment+* neurons responding to regression of above velocity measures against firing (N = 69 neurons). ***L,*** PCA projection of the regression measures, coloured according to significance of regression. ***M,*** Contribution of the regression measures towards the PC1 dimension. ***N,*** Contribution of the regression measures towards the PC2 dimension.

### Symmetrical feedback inhibition in VMHvl

Subsequently, we took advantage of the optrode design which allows blue light to be delivered in the vicinity of the recording electrodes to measure evoked electrophysiological responses in animals in which the light-sensitive cationic channel, Channelrhodopsin (ChR2), was expressed in specific VMHvl neuron populations. Previous *in vitro* electrophysiological and anatomical studies provided some evidence for bidirectional synaptic connectivity between inhibitory neurons in the VMHvl shell and core excitatory neurons (Yamamoto et al., 2018; Hashikawa et al., 2017) as well as for local connections between excitatory core neurons (Kennedy et al., 2020; Shao et al., 2022; **Figure 4A**). These data formed the basis for a computational model of VMH that explained the abrupt reciprocal firing changes of *Assessment*+ and *Flight*+ neurons as a nonlinear thresholding of a ramping input to VMH (Rahy et al., 2022). Exploration of optimal model parameters *in silico* showed that asymmetric feedback inhibition was necessary to impart nonlinear thresholding to the model. Specifically, the thresholded reciprocal switch in firing of *Assessment*+ and *Flight*+ neurons required either higher feedback inhibition on core *Assessment*+ neurons or a paradoxical, slow metabotropic excitatory component of reciprocal feedback inhibition onto *Flight*+ neurons (Rahy et al., 2022).

**Figure 4.**
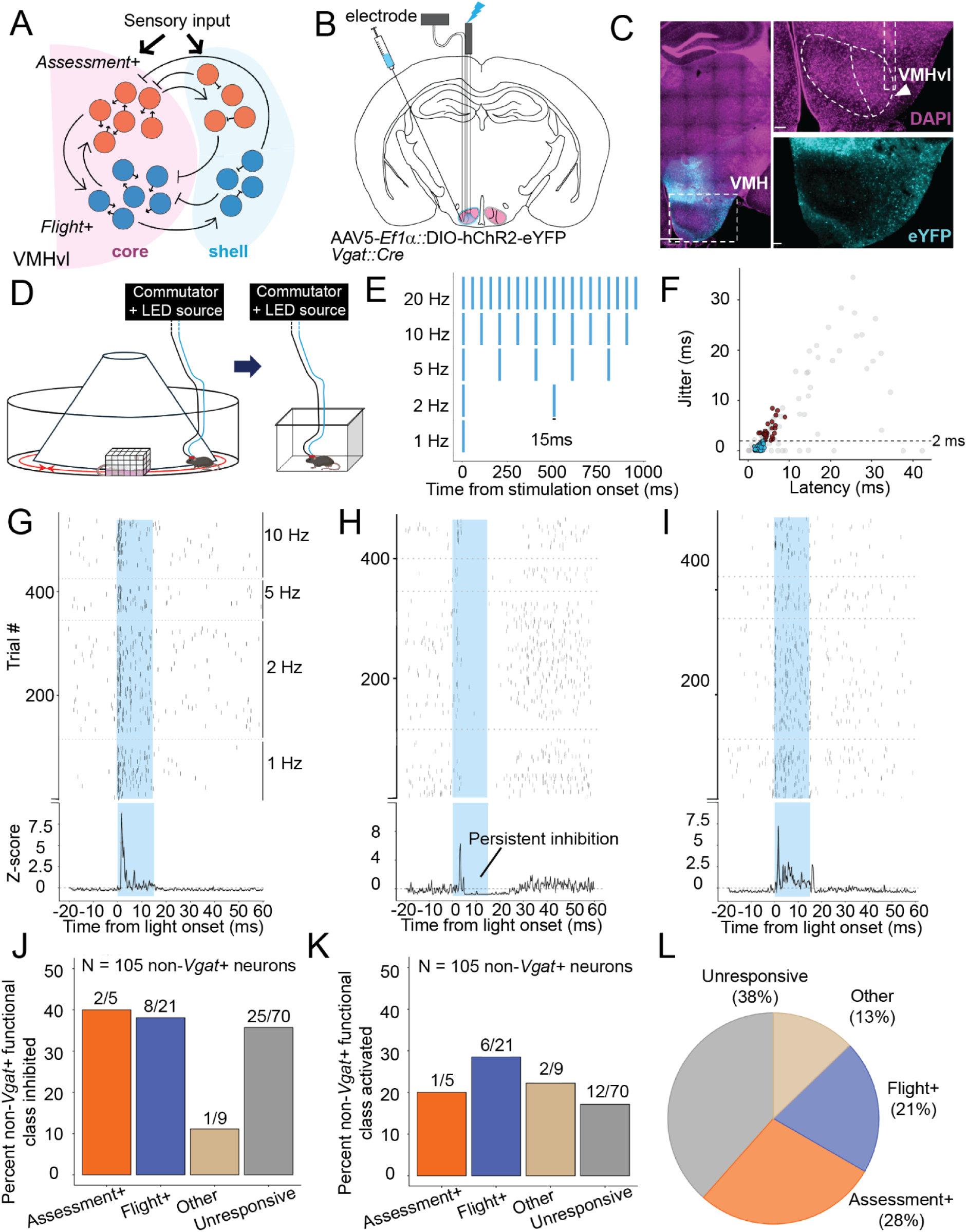
Characterization of VMHvl shell-to-core connectivity. ***A***, Proposed model depicting the connectivity among *Assessment+* (orange) and *Flight+* (blue) neurons of the VMHvl core (pink) and shell (light blue). ***B***, Viral strategy for opto-tagging and recording from the VMHvl shell neurons. ***C***, (**left**) Histology showing the expression of channelrhodopsin in shell (scale bar = 500 µm); (**right top**) DAPI staining showing the nuclei of neurons and outlines of the VMH with the ventrolateral subdivision and recording tract; (**right bottom**) eYFP expression indicating expression primarily in the shell and not core of VMHvl (scale bar = 100 µm). ***D***, Cartoon of experimental mouse undergoing light stimulation. ***E***, Stimulation frequencies used to check for light-responses within the nucleus. Each pulse was 15 ms long with an inter-pulse interval decided by the frequency of stimulation. ***F***, Plot of latency vs jitter of the first spike response of neurons within 49 s of light onset (N = 144 neurons). The neurons that were activated in the first 6 ms after light onset (N = 64 neurons) were divided into those with less (somatic response, blue, N = 39) or more (synaptic response, brown, N = 25) than 2 ms jitter. The remaining neurons (grey, N = 80) were determined to be either inhibited or unresponsive to light stimulation. ***G,*** Representative example of a low latency and low jitter somatic response by a neuron putatively expressing channelrhodopsin. (**top**) Raster plot of the firing of the neuron across light pulses of the different stimulation frequencies, separated by dashed lines (from bottom: 1, 2, 5, 10 Hz, bin width = 0.2 ms). (**bottom**) Average response PSTH across all trials. ***H,*** Representative example of an inhibitory response by a neuron likely receiving synaptic input from a channelrhodopsin expressing neuron. (**top**) Raster plot of the firing of the neuron across light pulses of the different stimulation frequencies, separated by dashed lines. (**bottom**) Average response PSTH across all trials. ***I,*** Representative example of a paradoxical, activatory response by a neuron. (**top**) Raster plot of the firing of the neuron across light pulses of the different stimulation frequencies, separated by dashed lines. (**bottom**) Average response PSTH across all trials. ***J,*** Distribution of the non-*Vgat+*, putative core neurons receiving synaptic inhibition in the post-stimulation period (16-25 ms after light onset) across the behavioral classes (non-*Vgat+* neurons: N = 105 neurons). Pairwise Fisher’s test: *Assessment+* vs *Flight+ p_adj_ = 0.93*, *Assessment+* vs *Others/Unresponsive p_adj_ = 0.74*, *Flight+* vs *Others/Unresponsive p_adj_ = 0.65*. ***K,*** Distribution of the non-*Vgat+* neurons showing synaptic activation in the stimulation period (5-14 ms after light onset) across the behavioral classes. Pairwise Fisher’s test: *Assessment+* vs *Flight+ p_adj_ = 0.69*, *Assessment+* vs *Others/Unresponsive p_adj_ = 0.89*, *Flight+* vs *Others/Unresponsive p_adj_ = 0.28*. ***L,*** Distribution of *Vgat*+ neurons across the behavioral classes (neurons with somatic response: N = 39 neurons, total neurons recorded: N = 144 neurons). *Assessment+* = 11/39, *Flight+* = 8/39, *Others* = 5/39, *Behaviorally unresponsive* = 15/39. Pairwise Fisher’s test: *Assessment+* vs *Flight+ p_adj_ = 0.02*, *Assessment+* vs *Others/Unresponsive p_adj_ < 0.001*, *Flight+* vs *Others/Unresponsive p_adj_ = 0.45*.

To confirm the existence of feedback inhibition and excitation in VMHvl *in vivo* and gather data for potential differential *Assessment*+ and *Flight*+ neuron connectivity, single unit recordings were performed in mice in which ChR2 was expressed in inhibitory VMHvl shell neurons (AAV-*Ef1a*::DIO-hChR2-eYFP, *Vgat*::Cre; **Figure 4B, C**). Evoked recordings were carried out in the home cage under quiet awake conditions at the end of the recording day (**Figure 4D**). Light pulses of increasing frequency (15 ms at 1, 2, 5, 10, 20 Hz; **Figure 4E**) were delivered to the surrounding tissue via the optrode, and the variation in latency (jitter) of evoked responses occurring in the first 6 ms after the 1 Hz stimulation were used to classify somatically activated vs. synaptically activated neurons (**Figure 4F**; see Materials & Methods; Lima et al., 2009). Forty-four percent (64/144) of neurons were rapidly activated in response to light stimulation (< 6 ms; **Figure 4G**) and were considered further for classification. Based on the jitter of the response across stimulation trials, 61% of these (jitter < 2 ms, 39/64) were classified as somatically activated ChR2-expressing *Vgat+* neurons (27% of all neurons, 39/144). Synaptic inhibition was generally characterized by a persistent suppression of firing following light delivery (**Figure 4H**). Thirty-four percent (36/105) of putative non-*Vgat*+ excitatory core neurons received significant inhibition, confirming the previously described shell-to-core inhibitory connectivity (Yamamoto et al., 2018; Minakuchi et al., 2024), while 54% (21/39) of *Vgat*+ neurons received significant inhibition, confirming prominent shell-to-shell connectivity (**Figure 4J, Figure S3B**). Unexpectedly, we also detected synaptic activation in 20% (21/105) of non-*Vgat*+ and 49% (19/39) of *Vgat*+ neurons (**Figure 4I, K**; **Figure S3C**). The presence of synaptic excitation upon the optogenetic activation of *Vgat*+ neurons suggests the presence of prominent multi-synaptic, double inhibitory connectivity in VMHvl.

To understand whether shell-to-core inhibition in VMHvl might be biased onto *Assessment*+ or *Flight*+ neurons, we compared light-evoked response patterns in the two behaviorally defined subclasses (**Figure S3A**). A similar fraction of putative core excitatory *Assessment*+ (40%, 2/5) and *Flight*+ (38%, 8/21) neurons showed significant inhibition following light stimulation, as did putative core excitatory neurons that did not show behavioral modulation (unresponsive, 36%, 25/70), arguing for an unbiased functional distribution of shell-to-core inhibitory inputs in VMHvl (**Figure 4J**). Similarly, no significant differences were seen in ChR2-evoked paradoxical excitation onto *Assessment*+ (20%, 1/5), *Flight*+ (29%, 6/21), or unresponsive neurons (17%, 12/70; **Figure 4K**).

### Assessment+ and Flight+ neurons in VMHvl shell

A major prediction of the VMH feedback inhibition model is the emergence of *Assessment*+ and *Flight*+ neuron properties among *Vgat*+ inhibitory shell neurons receiving inputs from excitatory core *Assessment*+ and *Flight*+ neurons, respectively (Rahy et al., 2022). To test this hypothesis, we examined the behavioral correlations of putative shell neurons (*Vgat*+ ChR2+). Cells with *Assessment*+ (28%, 11/39) and *Flight*+ (21%, 8/39) firing properties indistinguishable from those found in the VMHvl core were identified among *Vgat*+ ChR2+ neurons (**Figure 4L**) supporting a circuitry with prominent core-to-shell *Assessment*+- *Assessment*+ and *Flight*+-*Flight*+ connectivity. These data are consistent with the VMH feedback inhibition model (Rahy et al., 2022).

### Reciprocal feedback inhibition in VMHvl

The presence of excitatory feedback connections between core VMHvl neurons has been demonstrated *in vitro* (Kennedy et al., 2020; Shao et al., 2022). To confirm and characterize such connections *in vivo,* optrode recordings were carried out in the VMHvl of animals in which one of two dual-recombinase retrograde labelling approaches was used to express ChR2 exclusively in a subset of VMHvl core neurons that project to the dorsal periaqueductal grey (dPAG; strategy 1: AAVretro-*Ef1a*::DIO-FLPo in dPAG and AAV5-*CAG*::FLEXFRT-ChR2- mCherry in VMHvl of *Vglut2*::Cre mice; strategy 2: AAVretro-*CAG*::Cre in dPAG and AAV- *Ef1a*::DIO-hChR2-eYFP in VMHvl of C57/BL6J mice; **Figure 5A, B**). Following the classification of *Assessment+* and *Flight+* neurons during behavioral testing, the animals were returned to a holding cage and subjected to light-evoked optogenetic stimulation. As before, short latency responsive neurons (< 6 ms) were classified on the basis of jitter (< 2 ms, **Figure 5C**) to be somatically activated, core excitatory dPAG projection neurons (8%, 24/293, ChR2+, see example **Figure 5D**; see Materials & Methods). Significant synaptic activation was demonstrated in 13% (37/293, example **Figure 5E**) of neurons (both ChR2+ and ChR2–) and in nearly half of the cases (43%, 16/37) this activation persisted after light offset, confirming core-core excitatory connectivity and suggesting the existence of a metabotropic or other slow neurotransmission component to this excitation. Unexpectedly, we also observed a considerable number (20%, 58/293) of neurons receiving synaptic inhibition during light stimulation which in most cases consistently persisted after light offset (78%, 45/58; example **Figure 5F**). This response could reflect multi-synaptic feedback inhibition between core excitatory neurons via GABAergic shell neurons. Among excitatory VMHvl core neurons projecting to dPAG (ChR2+, somatically activated) an equal number were classified as *Assessment+* (25%, 6/24) or *Flight+* (25%, 6/24) neurons showing that both neuron classes contribute similarly to VMHvl projection neurons (**Figure S3E**). We observed no bias in either the excitatory synaptic input (feedback excitation, 19%, 6/31 vs. 4%, 1/25; **Figure S3F**) or the inhibitory synaptic input (feedback inhibition, 19%, 6/31 vs. 8%, 2/25; **Figure S3G**) onto ChR2- *Assessment+* and *Flight+* neurons.

**Figure 5.**
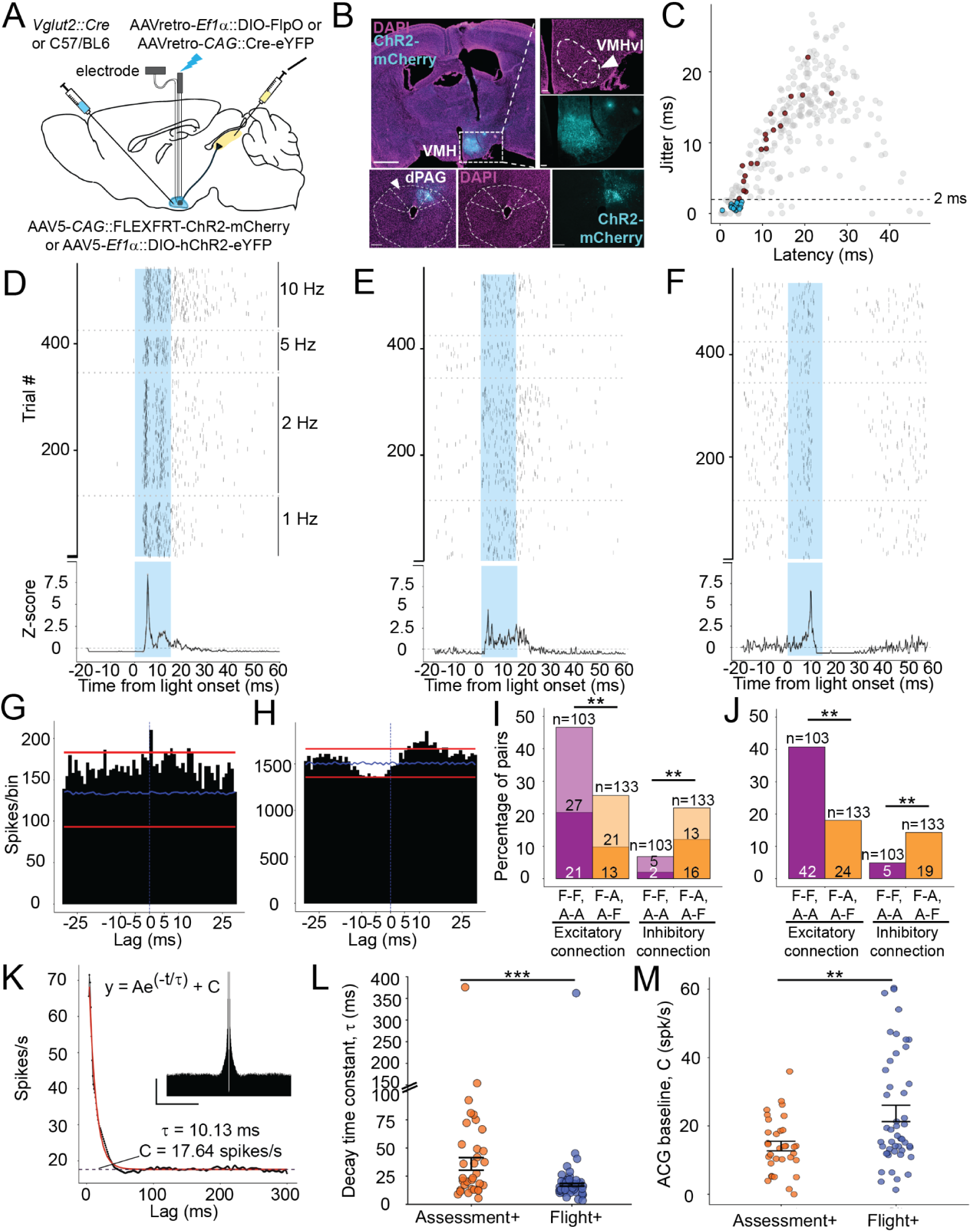
Characterization of VMHvl core-to-core connectivity. ***A***, Alternative viral strategies for opto-tagging and recording from dPAG-projecting neurons in VMHvl. The first strategy targets only dPAG-projecting, excitatory core neurons, while the second strategy labels all dPAG-projecting neurons. ***B***, (**first row, left**) Histology showing the expression of channelrhodopsin in *Vglut2+* VMH neurons (scale bar = 1 mm); (**first row, top right**) DAPI staining showing the nuclei of neurons and outline of the VMH; (**first row, bottom right**) mCherry expression in VMH confirming opsin expression (scale bar = 100 µm); (**bottom row, left**) histology showing the retrograde virus injection site in dPAG (scale bar = 800 µm); (**bottom row, middle**) DAPI staining showing outline of PAG; (**bottom row, right**) expression of mCherry marking the retrograde virus injection site. ***C***, Plot of latency vs jitter of the first spike response of neurons within 49 s of light onset (N = 293 neurons). Neurons that were activated in the first 6 ms after onset of light stimulation (N = 44 neurons) were divided into those with <2 ms jitter (somatic response, blue, N = 24) and those with > 2 ms jitter (synaptic response, brown, N = 20). The remaining neurons (grey, N = 249) were determined to be either inhibited or unresponsive to light stimulation. ***D***, Representative example of a low-latency and low-jitter, somatic response by a unit expressing channelrhodopsin. (**top**) Raster plot of the firing of the unit across light pulses of different stimulation frequencies, separated by dashed lines (from bottom: 1, 2, 5 and 10 Hz, bin width = 0.2 ms). (**bottom**) Average response PSTH across all trials. ***E***, Representative example of an excitatory response of a unit receiving feedback synaptic input. (**top**) Raster plot of the firing of the unit across light pulses of different stimulation frequencies, separated by dashed lines. (**bottom**) Average response PSTH across all trials. ***F***, Representative example of an inhibitory response of a unit receiving feedback synaptic input. (**top**) Raster plot of the firing of the unit across light pulses of different stimulation frequencies, separated by dashed lines. (**bottom**) Average response PSTH across all trials. ***G***, Representative example of a cross-correlogram between simultaneously recorded *Assessment+* and *Flight+* neurons showing fast onset, putatively monosynaptic, excitatory connectivity. ***H***, Representative example of a cross- correlogram between a pair of *Assessment+* and *Flight+* neurons showing fast onset, inhibitory connectivity (bin size = 1 ms). The blue line denotes bin-wise mean count/bin and the red lines indicate 1% and 99% confidence intervals derived from the null distribution. ***I***, Comparison of the percentage of like (*Assessment+-Assessment+* or *Flight+-Flight+*, purple, N = 103 pairs) and unlike (*Assessment+-Flight+* or *Flight+- Assessment+*, yellow, N = 133 pairs) neuron pairs showing excitatory or inhibitory connectivity. Numbers of A-A connections among like pairs and A-F connections among unlike pairs are indicated by the darker portion and F-F connections among like pairs, and F-A connections among unlike pairs by the lighter portion (Chi-square test of independence for excitatory connections: like pairs vs unlike pairs *p = 0.002*; Fisher test of independence for inhibitory connections: like pairs vs unlike pairs *p = 0.002*)*. **J***, Percentage of like and unlike pairs receiving common excitatory or inhibitory inputs (Chi-square test of independence for common excitatory input: like pairs vs unlike pairs *p < 0.001*; Fisher test of independence for common inhibitory input: like pairs vs unlike pairs *p = 0.017*)*. **K***, Representative example of mathematical fitting of the decay of the autocorrelogram. Depicted are the high-pass filtered autocorrelogram histogram (in black) and the predicted fit (in red) according to the exponential equation. The decay time constant τ and baseline of the autocorrelogram C (dashed black line) are represented. (**inset**) Autocorrelogram at ±300 ms lag (bin size = 1 ms, scale bar: x-axis = 100 ms, y-axis = 1000 spikes/bin). ***L***, Comparison of the decay time constant τ of neurons of the functional classes modelled with the exponential decay (*Assessment+*: N = 32/69 vs *Flight+*: N = 45/74; Mann-Whitney U test, W = 1095, *p = 0.001*; Fligner-Killeen test for homogeneity of variances, χ^2^(1) = 17.38, *p < 0.001*; Error bars represent mean ± SEM). ***M***, Distribution of the autocorrelogram baseline C of the neurons of the functional classes (Mann-Whitney U test, W = 510, *p = 0.009*; Error bars represent mean ± SEM; * *p < 0.05, ** p < 0.01, *** p < 0.001*).

Lastly, we sought evidence for a potential exclusivity of reciprocal feedback inhibition and excitation between excitatory core *Assessment+* and *Flight+* neurons. If feedback inhibition were exclusive, then individual *Assessment+* neurons should exhibit feedback inhibition from *Flight+* neurons, but not from other *Assessment+* neurons, and vice versa. Such connectivity can be inferred by measuring cross-correlations between pairs of connected neurons recorded simultaneously. Out of a total of 675 neurons recorded in our study, 236 neuron pair combinations were assessed from experimental sessions when *Assessment+* and *Flight+* neurons were recorded simultaneously. Statistical analysis of cross-correlograms between these pairs identified 69 pairs with putative monosynaptic (< 5 ms lag) excitatory (see example: **Figure 5G**) or inhibitory (see example: **Figure 5H**) connections. Other 49 pairs showed putative polysynaptic or metabotropic (> 5 ms lag) excitatory or inhibitory connections (see example: **Figure S4A, B**). Finally, additional pairs showed symmetrical cross-correlograms that indicated either common excitatory (see example: **Figure S4C**) or inhibitory (see example: **Figure S4D**) inputs to both neurons. Classifying the connected pairs (**Table S1**) revealed a non-random distribution of connectivity between *Assessment+* and *Flight+* neurons. While like pairs (*Assessment+-Assessment+*, A-A or *Flight+-Flight+*, F-F) showed a greater propensity to form putative excitatory connections than unlike pairs (*Flight+-Assessment+*, F-A or *Assessment+-Flight+*, A-F; 47%, 48/103 vs 26%, 34/133, respectively), the reverse was true for inhibitory connections (7%, 7/103 vs 22%, 29/133; **Figure 5I, Table S1**) demonstrating a strong enrichment of heterotypic (A-F or F-A) feedback inhibition.

Moreover, common feedback excitation was significantly more frequent in like (41%, 42/103) than unlike (vs. 18%, 24/133) pairs (**Figure 5J**) demonstrating a bias that favours excitatory connections between neuron classes. Surprisingly, common inhibitory inputs were also depleted for like pairs (A-A or F-F; 5%, 5/103) compared to unlike pairs (A-F or F-A; 14%, 19/133; **Figure 5J**). The reason for the depletion of common inhibitory inputs in like pairs is not clear at present, but may simply reflect a difficulty in detecting common inhibition in pairs that are receiving common excitation coupled to the higher probability of common excitation in like neuron pairs. The bias in the common inputs resembles the pattern of the respective putative connections, indicating that one probable source of these inputs might be the synaptic connections within the nucleus spreading the feedback between pairs of neurons. In summary, the cross-correlation data confirm the existence of feedback excitation between VMHvl neurons with a greater propensity for this to occur within *Assessment+* or *Flight+* neuron populations, and provide strong support for the existence of selective reciprocal feedback inhibition between *Assessment+* and *Flight+* neurons as predicted by the computational model (Rahy et al., 2022).

### Differences in the intrinsic firing properties of Assessment+ and Flight+

Although we did not find evidence for significant differences in synaptic connectivity between the functional classes as predicted by the computational model (Rahy et al., 2022), we did find evidence for possible differences in intrinsic firing properties between *Assessment*+ and *Flight*+ neurons, a feature that had not been considered in the computational model, but that has been reported in earlier *in vitro* and *in vivo* anesthetized VMH recordings (Sabatier and Leng, 2008; Yamamoto et al., 2018; MacGregor and Leng, 2019). While all *Assessment+* ChR2+ neurons (100%, 6/6) were persistently activated during somatic light stimulation (**Figure S3K, M**; **Figure S4I, K**), the majority of *Flight+* ChR2+ neurons (83%, 5/6) were transiently activated and then showed an inhibition of firing (**Figure S3M, N**; **Figure S4J, K**). Notably, this difference in evoked response pattern was not seen among the respective ChR2- groups, suggesting that it might reflect differences in intrinsic firing properties rather than any bias in connectivity. To confirm this hypothesis, we inspected the autocorrelograms of *Assessment+* and *Flight+* neurons to assess the temporal structure of a neuron’s firing. The decay in autocorrelograms across time lags were fitted with an exponential function whenever possible (80/143 neurons, *Assessment+* = 33/69, *Flight+* = 47/74) and the decay time constant, τ, extracted (**Figure 5K**). Consistent with the difference in firing responses shown to somatic optrode stimulation, *Flight+* neurons showed a significantly lower decay time constant (24.3 ± 7.4 ms) when compared to *Assessment+* neurons (48.13 ± 11.16 ms), which also showed a significantly wider distribution of the time constants (**Figure 5L**). Moreover, a greater fraction of *Flight+* neurons had short decay time constants (τ ≤ 30 ms, 58%, 43/74) compared to *Assessment+* neurons (26%, 18/69; Chi-square test: *p < 0.001*), suggesting increased burst-like firing responses in *Flight+* neurons. Notably, these differences were confirmed both among the VMHvl core (**Figure S4F**; τ ≤ 30 ms, *Assessment+* vs. *Flight+*: 0/11, 0% vs. 20/27, 74%; Fisher test: *p < 0.001*) and shell (**Figure S4H**; τ ≤ 30 ms, *Assessment+* vs *Flight+*: 1/11, 9% vs 5/8, 63%; Fisher test: *p = 0.04*). Additionally, *Flight+* autocorrelograms decayed to a significantly higher baseline firing rate than *Assessment+* neurons (23.7 ± 2.4 spikes/s vs 14.1 ± 1.4 spikes/s in 300 ms lag; **Figure 5M**), with neurons of both classes that exhibited a faster decay showing higher baseline firing (**Figure S4E**). Finally, we compared the coefficient of variation (CV) to assess regularity in firing patterns of the *Assessment+* and *Flight+* neurons. CV of the functional classes were extracted from the respective behavioral epochs when they showed increased firing, during the two seconds period before social investigation offset for *Assessment+* neurons and during the two seconds period after flight onset for *Flight+* neurons. During these periods the *Assessment+* population showed significantly higher CV values, hence more irregular firing, than *Flight+* neurons (**Figure S4L**). Taken together, we found that *Flight+* neurons showed a faster modulation of both their evoked and spontaneous firing responses than *Assessment+* neurons, demonstrating significant differences in the intrinsic firing properties of these neuron classes and revealing a potential mechanism to explain the asymmetry of their firing responses during the approach-to-escape transition.

## Discussion

In this study, we used single-unit optrode electrophysiology in the VMHvl of awake behaving mice to 1) investigate the neural encoding of approach-escape behavior toward a social threat, and 2) test specific features of local neural connectivity predicted to explain that neural encoding. Our experiments were guided by previous studies that identified two major classes of neurons in the VMH, *Assessment+* and *Flight+* neurons (Esteban Masferrer et al., 2020), and by a computational model of VMH that made specific predictions about local neural connectivity among *Assessment+* and *Flight+* neurons (Rahy et al., 2022). Our findings confirm the presence of *Assessment+* and *Flight+* neurons in VMHvl previously reported using *in vivo* calcium imaging (Krzywkowski et al., 2020). We also confirmed three predictions of the computational model: 1) the existence of *Assessment+* and *Flight+* neurons amongst GABAergic neurons surrounding the VMHvl core, known as the VMHvl shell, 2), the presence of feedback excitation between VMHvl core neurons, and 3) the presence of reciprocal feedback inhibition between VMHvl core neurons (**Figure 6**). However, we were not able to confirm a fourth prediction of the computational model, namely, an asymmetry in the feedback inhibition between *Assessment+* and *Flight+* neurons. Instead, we discovered differences in the intrinsic firing properties of *Assessment+* and *Flight+* neurons, leading us to predict that these intrinsic differences may impart the asymmetry required by the computational model for the emergence of *Assessment+* and *Flight+* neuron classes.

**Figure 6.**
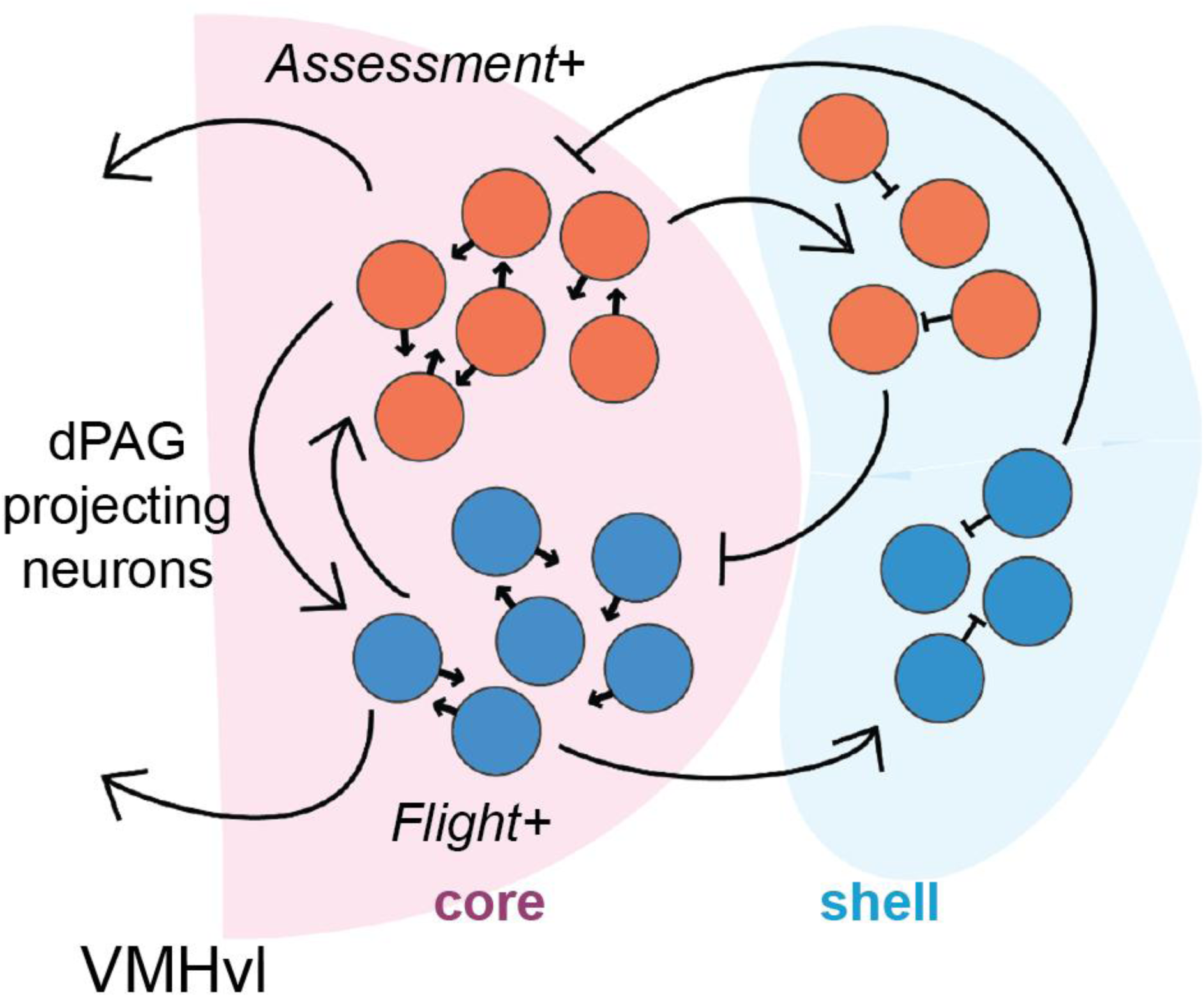
Circuit model of VMHvl. Model of local circuitry of VMHvl depicting the connectivity among *Assessment+* (orange) and *Flight+* (blue) neurons in core (pink) and shell (light blue) based on results from this and other studies. Note that both *Assessment+* and *Flight+* neurons project to dPAG and darker arrows indicate greater connectivity within a functional class.

An important advance of our study was the development of a compact circular behavioral apparatus compatible with tethered neural recording that allows pairs of mice to freely pursue each other until the chase comes to its natural conclusion (**Figure 1A-B, Figure S1A-D**). This feature allowed us to carry out *in vivo* single-unit electrophysiological recordings to assess neural correlations with the full range of escape behaviors from start to conclusion in a manner not possible with previous apparatuses (Lin et al., 2011; Silva et al., 2013). Nevertheless, we found that the relatively small circular design needed to allow tethered *in vivo* recording required animals to continuously reorient their pursuit behavior and was associated with decreased peak escape velocities when compared to apparatuses allowing less-constrained trajectories (Esteban Masferrer et al., 2020). Social pursuit in larger semi-natural arenas do achieve high peak velocities (Battivelli et al., 2024), suggesting that an enlargement of the apparatus could be considered to ameliorate this limitation in the future.

Previous studies using *in vivo* calcium miniscopes had identified putative *Assessment+* and *Flight+* neurons in VMHvl (Krzywkowski et al., 2020). However, the poor temporal resolution and propensity for motion artifacts associated with miniscope calcium imaging left it unclear whether these neurons had properties similar to those described in the nearby VMHdm during approach-escape from a live rat (Esteban Masferrer et al., 2020). Our findings revealed *Assessment+* and *Flight+* neuron firing patterns in VMHvl to be highly similar to those found in the VMHdm (Esteban Masferrer et al., 2020), with an abrupt switch in firing activity at the escape onset point (**Figure 2E-F**, see also **Figure S2F-I**). Notably, in the few cases in which *Assessment+* and *Flight+* neurons were recorded simultaneously in the same animal and session, their firing showed a precise inverse correlation with *Assessment+* neurons shutting off their firing exactly at the moment in which *Flight+* neurons increased theirs (**Figure 2C-D**). Moreover, a trial-by-trial analysis of the behavioral correlates of this firing switch showed it to coincide or directly precede the increase in velocity associated with the escape bout (**Figure S2H-J**). Importantly, the magnitude of the increase in *Flight+* neurons firing at escape onset correlated across trials with the increase in velocity (**Figure 3C-D**) consistent with a role in setting the vigor of escape behavior. This was not the case for *Assessment+* neurons (**Figure 3J-K**). As a result, we argue that *Assessment+* neurons are more likely to be involved in setting the probability of escape, while *Flight+* neurons could have a role in setting its quality. This interpretation is consistent with the ability of optogenetic activation of VMHvl core neurons to elicit avoidance behavior (Sakurai et al., 2016; Wang et al., 2019; Krzywkowski et al., 2020).

Our data show that *Assessment+* and *Flight+* neurons are a common feature of the medial hypothalamic nuclei controlling defensive behavior regardless of the nature of the threat (Canteras, 2002; Gross and Canteras, 2012) and suggest a common computational role for these nodes in driving behavior. The findings are also important because previous single-unit recording studies in VMHvl reported *Attack+* neurons (Lin et al., 2011; Falkner et al., 2014; Lee et al., 2014) that increased their firing during approach-attack behavior, but left open the question as to whether approach-escape transitions could similarly be encoded in this structure. Although we did not record during conspecific aggression in this study, we speculate that *Assessment+* neurons at least partially overlap with *Attack+* neurons because they both increase their firing during social investigation (Lin et al., 2011). Consistent with this hypothesis, *in vivo* single-unit calcium imaging data collected from VMHvl during attack and escape using miniscopes found that more than a third of *Attack+* neurons had *Assessment+* properties as assessed when the animals were later subjected to social defeat, and cFos serial-tagging experiments found similar overlap between aggression and defeat (Krzywkowski et al., 2020).

A major goal of the study was to test several predictions emerging from a circuit model of VMH that attempted to explain the firing patterns of *Assessment+* and *Flight+* neurons during approach-escape trials (Rahy et al., 2022). The model assumed the presence of excitatory *Assessment+* and *Flight+* neurons in the VMH core receiving excitatory sensory input from upstream areas carrying threat information that are interconnected via feedback excitation and inhibition. An unbiased exploration of connectivity parameters in the model identified conditions that resulted in *Assessment+* neurons ramping up their activity during approach while *Flight+* neurons remained silent, followed by the spontaneous triggering of a switch in activity in which *Assessment+* neurons silenced their firing and *Flight+* neurons were activated. Not unexpectedly, conditions under which this nonlinear switch occurred *in silico* all required asymmetry in the synaptic connectivity between *Assessment+* and *Flight+* neurons. However, while our experimental data confirmed the presence of feedback excitation (**Figure S3F, I** and **Figure 4K**) and reciprocal feedback inhibition (**Figure 5I** and **Figure S3G, J**) as well as the presence of *Assessment+* and *Flight+* neurons among GABAergic VMHvl shell neurons (**Figure 4L**), they failed to detect any asymmetry in this connectivity. Instead, we uncovered a significant bias in the intrinsic firing properties of *Assessment+* and *Flight+* neurons, a difference reflected in both spontaneous and evoked firing activities (**Figure 5L-M** and **Figure S4I-K**). We speculate that this difference could give rise to the asymmetry between neuron classes required by the model for the emergence of the reciprocal switch in firing. If so, this raises the possibility that the switch in *Assessment+* and *Flight+* neuron firing at escape onset could depend entirely on local VMH circuit properties. We note that previous single-unit *in vitro* slice and *in vivo* anesthetized recordings in VMH have reported at least two major neuron classes with different intrinsic firing properties (Sabatier and Leng, 2008; Yamamoto et al., 2018; MacGregor and Leng, 2019).

In contrast, we also noted several unexplained features of our *in vivo* optrode recordings. For example, while neurons in the VMHvl core showed the expected inhibition after optogenetic stimulation of GABAergic shell neurons (**Figure 4H, J**), they also presented paradoxical activation (**Figure 4I, K**). The source of this activation is unclear at present, but was most prominent during the late period of light stimulation (5-14 ms; **Figure S3D**) suggesting a slow, multi-synaptic or metabotropic mechanism. Nuclei of the medial hypothalamus are highly reciprocally interconnected (Canteras et al., 1994) and the VMHvl shell has been shown to express neuropeptides, including somatostatin (Kim et al., 2019), features that could contribute to slow excitatory synaptic effects. Similarly, a combination of inhibition and activation was also seen in VMHvl shell-shell connections (**Figure S3B, C**), but it remained unclear if this was the result of the expected somatic ChR2-evoked excitation or a possible additional synaptic excitation phenomenon. Finally, our retrograde labeling experiment showed that *Assessment+* and *Flight+* neurons were equally likely to project to dPAG (**Figure S3E**). This balanced output has implications for how downstream target areas responsible for triggering the escape response operate, and suggests that *Assessment+* neuron outputs also have a role in promoting defense, potentially by promoting risk assessment behaviors, such as stretch-approach, stretch-attend, or immobility before the onset of escape. Distinguishing the functional contributions of the two neuron classes would require identifying genetic markers that would allow for their selective manipulation. Our discovery of differences in intrinsic firing properties between *Assessment+* and *Flight+* neurons offers a way forward and suggests that genes responsible for these differential biophysical properties could be exploited as genetic markers. Overall, our findings contribute to understanding how local microcircuits can threshold sensory inputs to produce innate defensive behaviors, with implications for the production of aversive emotional states produced by these nuclei in humans (Parvizi et al., 2022).

## Materials and Methods

### Animals

All experimental procedures involving the use of animals were carried out in accordance with the European Union (EU) Directive 2010/63/EU and according to protocols approved by the EMBL IACUC (2021-04-09_CG_amend7) and Italian Ministry of Health (183/2024-PR). Adult male C57BL/6J heterozygous *Vgat*::ires-Cre (Jax stock no. 028862; Vong et al., 2011) and heterozygous *Vglut2*::ires-Cre (Jax stock no. 028863; Vong et al., 2011) were bred in the EMBL Rome Laboratory Animal Resources facility. Animals were singly-housed after surgery under a 12-hour light-dark cycle and had access to food and water *ad libitum*. All experiments were performed in the light phase. Singly-housed CD1 ex-breeder males (Charles River Laboratories) were selected as aggressive interaction partners (aggressors) after a three-day resident-intruder screening (Franklin et al., 2017). Group-housed male CD1s bred in the EMBL Rome animal facility were used as non-aggressive interaction partners.

### Surgery

Surgeries were performed on 8 to 12-week-old male mice under deep anaesthesia (3-4% isofluorane in O_2_ for induction and 1-2% isofluorane for maintenance). Stereotaxic surgeries were performed on RWD Life Sciences or Kopf Instrument frames. For extra-cellular recordings coupled with optogenetic stimulation mice were implanted with moveable optrodes into the VMHvl (AP = -0.92 mm, ML = -0.65 mm, and DV = -5.80 mm with respect to bregma). Optrodes were constructed by attaching a 16-channel electrode and an optic fiber (200 µm core diameter, 0.5 numerical aperture, ceramic ferrule: 1.25 mm outer diameter, RWD Life Sciences) on a nano-drive (Cambridge Neurotech). The electrodes were either commercially available 16-channel silicon probes (9 mm E-2 probes, Cambridge Neurotech) or made by bundling 16 insulated tungsten microwires with bare tips (23 µm in diameter, impedance ∼300 kΩ, California Fine Wire) and a 300 µm diameter silver wire for grounding. Viruses and plasmids were either purchased from Addgene or produced by the EMBL Rome Gene Editing and Virus Facility (GEVF). For optogenetic activation of local inhibitory neurons, AAV5- *Ef1a*::DIO-ChR2(E123T/T159C)-eYFP (#35509, Addgene, Mattis et al., 2011) virus was injected into the VMHvl (AP = -0.92 mm, ML = -0.65 mm, and DV = -5.95 mm with respect to bregma) of *Vgat*::Cre mice. For optogenetic activation of excitatory neurons, AAV5- *Ef1a*::DIO-ChR2(E123T/T159C)-eYFP virus was injected into the VMHvl of *Vglut2*::Cre mice. For the optogenetic activation of PAG-projecting VMHvl neurons AAV5-*Ef1a*::DIO- ChR2(E123T/T159C)-eYFP or AAV5-*CAG*::FLEX-FRT-ChR2-mCherry (#75470, Addgene, Tang et al., 2016) were injected into the VMHvl, while AAVretro-*CAG*::iCre-T2A-eGFP (GEVF) or AAVretro-*Ef1a::*DIO-FlpO (#87306, Addgene, Zingg et al., 2017) was injected into the dPAG (AP = -4.0 mm, ML = -1.56 mm, and DV = -2.35 mm with respect to bregma, at 26° lateral angle) of wild-type or *Vglut2*::Cre mice, respectively. Three stainless steel screws (RWD Life Sciences) were fixed permanently into the posterior and anterior portions of the skull to serve as ground and fixation. Implants were fixed on the skull using dental acrylic (DuraLay Reliance). After surgery mice were given analgesia in drinking water (7.6 mg/10 mL paracetamol) for 7 days to control pain and inflammation. All animals were allowed to recover for at least two weeks before behavioral habituation and testing.

### Social pursuit test

A new behavioral test was developed by adapting the previous repeated social interaction test (Krzywkowski et al., 2020) to include free social interaction between the experimental mouse and social aggressor. The test arena was modified into a circular corridor of ∼172 cm circumference (60 cm outer diameter, 5 cm wide) constructed with outer walls of 30 cm height in transparent plastic and inner walls formed by a cone (∼50 cm height and 40 cm diameter). The base of the arena was covered with a yellow plastic sheet to allow for video tracking of the black-coated experimental mouse as well as white-coated social aggressor. Each session consisted of five phases. Behavioral testing commenced with a 5 min Initial Exploration of the arena with a wiremesh cage placed in the designated social zone of the corridor. A screened aggressive CD1 or a non-screened, group-housed CD1 was introduced into the wiremesh cage in the successive Restricted Interaction (RI) phase, which ran for a period of 10 min or until the mouse completed 10 assessment-escape trials. Subsequently, the cage was removed and the mice allowed to interact freely for 10 min in the Free Interaction (FI) phase. The phase was occasionally ended early to prevent bodily harm if attacks by the aggressive mouse were judged to be too strong, or the experimental mouse behaved in a manner consistent with being strongly defeated. In order to maintain a level of social fear that facilitated assessment-escape behavior, the social threat was chosen to either be an aggressive CD1 or a non-screened counterpart. The choice was based on the intensity of defeat in the previous session, time between behavioral exposure, and by gauging the exploratory and social avoidance behavior shown by the test mouse. In the following Restricted Interaction 2 (RI2) phase the conspecific was returned to the wiremesh cage in the social zone and the test mouse carried out assessment-escape trials for 10 min. The conspecific was removed and the wiremesh cage replaced for 5 min in the Post Exploration 1 phase, followed by removal of the cage for 5 min in the final Post Exploration 2 phase. A period of 24 hours was kept between exposures which was extended if the social defeat was gauged to be too strong based on low mobility/freezing or very low exploratory behavior of the mice in the phases after FI.

Behavior was recorded at 50 fps from the top and the side with two cameras using CinePlex Studio (Cineplex Behavioral Research System, Plexon) software. Frame-by-frame annotation was carried out manually using Solomon Coder (RRID:SCR_016041) or CinePlex Editor. The following behaviors were scored: *approach* – locomotion by the experimental mouse towards the caged/free threat; *investigation* – close contact with nose oriented towards the threat by the experimental mouse; *stretch attend* posture or *stretch approach*; *flight* – whole body movement away from the caged/free threat by the experimental mouse with locomotion over two body-lengths; *flight with pursuit* – rapid whole body movement away from freely moving threat by the experimental mouse along with pursuit by the threat, which occurred with or without attacks on the hind quarters of the experimental mouse; *short flight* – whole body movement or sudden retraction of body/head by the test mouse from the caged/free threat with slower locomotion or movement of less than 2 body-lengths; *defensive upright* – full or half rearing on hind paws by the experimental mouse with the front paws off the ground during close interaction with the freely-moving threat, often displaying its ventral side; *aggressor approach* – approach by the freely moving threat towards the experimental mouse; *attack* – attack by the experimental mouse with/without pursuit of the threat. Marker-less tracking of the body parts of the mice was carried out using DeepLabCut (Nath et al., 2019) with separate single-animal models for the experimental mouse and aggressor. The following sixteen body parts were tracked – nose, head, ears, neck, front limbs, center, sides, back, hind limbs, tail base, tail center and tail end. SimBA (Nilsson et al., 2020) was used to perform body part extrapolation and outlier correction of the position data. Any animals with misplaced viral infections or optic fiber implants were excluded from the analysis.

The frame-by-frame x and y coordinates of the centers of the animals were used to find the azimuth angle. The coordinates of the center of the arena were used to re-align the animals’ positions by subtracting it from the animals’ positions and the arc-tangent angles calculated, which ranged between [0, 359.9] degrees. In the RI and RI2 phases, the azimuth angle of the center point of the wiremesh cage was also calculated. The distance between the experimental mouse and the reference object (wiremesh cage or social threat) was calculated by finding a diagonal line dividing the arena symmetrically about the center of reference (defined as 0 degree), adjusting the azimuth of the experimental mouse with respect to the new 0 degree, and calculating the angle difference which ranged between [0, 180] degrees. The percent time spent around the reference object in the RI, FI and RI2 phases was calculated by finding the duration when the center of the experimental mouse was under 60-degree angle away from the center of reference object with respect to the duration of the behavioral phase.

### In vivo electrophysiology

During the recording sessions mice were connected to a lightweight head stage (1.03 g, gain 20×, Plexon). The head stage was connected to a 16-channel analog amplifier (gain 50×, Plexon) and neural activity was checked online. The common median or common average signal of the channels were used for referencing. If at least one unit was identified, the behavioral paradigm was started. Otherwise, the bundle of electrodes was re-adjusted by moving the screw to advance it by 62.5 µm followed by a waiting period of 24 hours before the next recording session. The neural signal was acquired (digitized at 40 kHz) and filtered to separate the high frequency signal (300 Hz to 8 kHz) using a Neural Data Acquisition System (Omniplex, Plexon). Information was stored for offline analysis. Spikes were sorted offline from the high frequency filtered data on Kilosort2 GUI (Pachitariu et al., 2024) using custom made channel maps- either based on the geometry of the silicon probe or a linear map for the wire electrodes made in the lab, followed by manual annotation and curation of unit clusters on Phy2 (Rossant et al., 2016). Unit isolation was verified using autocorrelation histograms, coherence of waveforms and principal component analysis (PCA) projection. Cross-correlation histograms were used to detect neurons appearing in more than one channel. VMHvl neurons were recorded from 17 mice across 89 sessions. Neurons with < 0.1% percentage of spikes occurring with ISIs < 1.2ms were deemed to be single-units (262/676; Nomoto and Lima, 2015), those with > 0.1% and < 2% of spikes within ISI were deemed to be multi-units (413/676), and the rest discarded (1/676).

### Optogenetic-assisted circuit-mapping

Experimental mice were placed in a novel cage and optogenetic stimulation with a LED source (300 mA, 465 nm, 9-18 mW at the tip of the optic fibre) was delivered via high performance patch cables (Plexbright Optogenetic Stimulation System, Plexon). Stimulation frequencies (1, 2, 5, 10, 20 Hz) were used with a square pulse of 15 ms duration, or 1 Hz stimulation of 2 ms duration. In the instances where light-evoked artefacts were observed at the pulse onset and offset, additional 1 Hz ramped stimulation trials were used, with ramped onset, holding period and ramped offset of 20 ms each or pulses with holding period and ramped offset of 5 ms each with varying duration of ramped onset such as 5 ms, 10 ms and 15 ms. For the lower (1, 2 Hz) frequencies, stimulation was carried out for 100 repetitions, while for the higher frequencies the inter-stimulus interval was 59 s and was repeated 10 times. For the ramped stimulations, 10 repetitions were performed.

### Spike analysis

For each neuron a mean spike density function was computed by applying a Gaussian kernel (kernel window = 3 samples) to the spike counts with bin size = 20 ms on NeuroExplorer (Nex Technologies), corresponding to σ = 25.5 ms. Z-scored peristimulus time histograms (PSTHs) and peristimulus rasters were computed using custom written scripts in R Statistical Software (R Core Team, 2025). For the mean PSTH the firing rate for each unit was averaged across trials per mouse. For the mean baseline PSTHs used for statistical comparison the firing rate was averaged across randomly sampled times in the RI1 and FI test phases of the same number as the flight behavior trials. The baseline-subtracted firing per bin in the RI1 and FI test phases was calculated by subtracting the average of firing of 100 randomly sampled 20 ms bins from the Gaussian-convolved firing rate. For visual representation the trial averages were smoothed with a Gaussian filter with a window length n = 5 samples and shape parameter, α = 2.5 (Van Boxtel, G.J.M., et al., 2021). To create the light stimulation aligned raster plots and PSTHs for each unit the firing rate was calculated using 0.2 ms bins for a period of -20 ms to 60 ms from light onset for all frequencies, except 20 Hz for which a period of -10 ms to 40 ms was used. For plotting, the trial averages were smoothed with a Gaussian filter with a window length n = 3 samples and shape parameter, α = 5 (Van Boxtel, G.J.M., et al., 2021). Autocorrelograms for all *Assessment+* and *Flight+* neurons were created with a bin size of 1 ms and maximum lag of 300 ms. Cross-correlograms for connectivity analysis were created for all unit pair combinations recorded on the experimental days when both *Assessment+* and *Flight+* unit were detected along with 750 shift-predictors using a randomization procedure for the period of the behavioral exposure with bin size of 1 ms and maximum lag of 30 ms. The exposure period was divided into 20 s long trial windows and the shift predictors created by aligning the firing in the trial windows of the reference unit with the firing of the target unit in a randomly selected trial window. A shift-predictor subtracted histogram was created by subtracting the bin-wise average of the 750 shift-predictor histograms from the cross-correlogram for each pair.

### Statistics

All of the statistical analyses were performed using custom written scripts in R (R Core Team, 2025; Wickham et al., 2019; Van Boxtel, G.J.M., et al., 2021; Barrett et al., 2025; Wilke, 2025; Ushey et al., 2025; Garnier et al., 2024; Shuangbin Xu et al., 2021; Scrucca et al., 2023) or Python. To depict the change in position of the mice over the duration of a representative trial, the azimuth of the center of the mice was plotted over time coloured according to the movement velocity. The time axis was represented along the radius of the circle with the starting time at the center and ending time of the trial at the circumference of the circle. The azimuth projection of the edges of the stationary wiremesh cage was used to find the breadth of the sector in the circular corridor it occupied. To plot the variation in distance between the experimental mouse and reference object across multiple assessment-escape trials a subset of all trials was randomly sampled and plotted along with the trendline predicted using generalised additive modelling (GAM). The difference in the total distance moved during the first 5 s and velocity changes during the first 2 s of a behavioral trial were measured by calculating the area under curve (AUC) of the velocity and acceleration traces respectively using the trapezoid method (Signorell, 2025). The average of all behavioral trials of a specific condition (e.g. flights in RI, flights in FI, and flights with pursuit in FI) from an experimental session was considered a datapoint for comparison and the distributions formed across sessions, across animals. Each distribution was checked for normality with a histogram and a quantile-quantile plot and compared with one-way ANOVA followed by a post-hoc Tukey HSD test to determine significance (*p_adj_ < 0.05*). To determine the difference in social interaction in the different test phases, the session-wise percent time spent close to the wiremesh cage (< 60 degree) was calculated while comparing the RI and RI2 phases and the time spent close to the social threat during investigation bouts (< 60 degree) while comparing RI and FI phases. The distributions were checked for normality as before and compared with paired Welch’s t-test or Wilcoxon signed rank test to check for significance (*p < 0.05*).

To identify neurons responsive to approach-escape switches the average firing in the 2 s pre-flight and 2 s post-flight periods were compared against average firing from one hundred 2 s long baselines with Mann-Whitney U test. Neurons were annotated as being behaviorally-responsive when the firing in either pre- or post-flight periods differed significantly from at least ninety baselines (*p < 0.001*). To identify clusters based on the pattern of activity the average PSTHs of the behaviorally-responsive neurons centered on the approach-escape transitions were clustered using a variant of density-based clustering (ordering points to identify the clustering structure, OPTICS; Hahsler et al., 2019). The average PSTH of each unit was down-sampled to 60 ms time bins and smoothed using a Gaussian filter with window of n = 3 samples, and shape parameter, α = 2.5 (Hamilton, 2004). A minimum of 5 neighbouring points was used to define a core point, and cutoff of ε = 8.35 applied after visual inspection of the ordered reachability distance plot to obtain two clusters along with unassigned noise neurons (**Figure S2A**). The clusters were annotated as *Assessment+* and *Flight+* based on their firing dynamics at flight onset while the noise cluster was annotated as Others. The average PSTH’s were normalized to have unit variance and mean of zero and projected onto PCA space for visualisation.

In order to evaluate the behavioral feature with which the escape switch coincided the start of movement was determined by aligning the PSTH’s of *Assessment+* and *Flight+* neurons. The moment of velocity peak was established in the period 2 s after flight onset and the time when the velocity increased to at least 20% of the peak velocity value was identified. To quantify the relationship between neural activity of *Flight+* and *Assessment+* neurons and locomotor behavior two complementary linear regression analyses were performed for each recorded unit in each mouse from behavior annotated in the RI1 and FI test phases. For *Flight+* neurons manually annotated flight behavior onsets were identified to serve as alignment points. For each behavioral trial: 1) the initial acceleration was determined as the velocity slope in a window spanning 200 ms before flight onset up to 40 ms after the first velocity peak within 2 s of onset, and the neural activity change as the slope of the baseline subtracted firing in the window from flight onset up to 40 ms after the velocity peak using linear regression; 2) the AUC of the velocity was calculated as a measure of displacement over the whole duration of the flight trial (Signorell, 2025), and the neural activity as the sum of the baseline subtracted firing in the 2 s after flight onset normalized by the window duration to obtain a spike-rate measure. For *Assessment+* neurons the approach-investigation transitions were identified to serve as alignment points. For each behavioral transition: 1) the initial acceleration was determined as the velocity slope in a window spanning 200 ms before behavioral transition up to minimum velocity within 0.5 s of the investigation onset and the neural activity change as the slope of the baseline subtracted firing in the same time window using linear regression; 2) the AUC of the velocity was calculated as a measure of displacement over the whole duration of the investigation trial (Signorell, 2025), and the neural activity as the sum of the baseline subtracted firing in the 2 s after investigation onset normalized by the window duration to obtain a spike-rate measure. The positional measures were regressed against the respective neural activity measures for each *Flight+* or *Assessment+* neuron and the regression slope (β), adjusted coefficient of determination (R²), and associated p-value were extracted. The p-values for the two regressions per unit were adjusted by using Benjamini-Hochberg method for multiple comparisons (*p_adj_ < 0.05*). PCA was used to linearly transform the output of the flight behavior aligned regressions after normalising the variable to have unit variance and mean of zero and project the *Flight+* neurons in a reduced dimensional space; similarly, PCA was calculated for the *Assessment+* neurons using the approach-investigation transition aligned regressions.

To qualify the recorded neurons as being responsive to optogenetic stimulation the total spikes in a certain time window after the onset of each light pulse were compared against an equal number of randomly selected windows of equal duration taken from the inter-pulse interval. The firing distributions were tested with Mann-Whitney U test (*p < 0.05*) and the light-responsive neurons annotated as activated or inhibited based on the difference between the average of the light-aligned and baseline responses. The latency and jitter of the unit’s response to light stimulation were calculated with latency being the median of the timing of the first spike detected within the initial 49 ms period after each light pulse onset and the jitter being the median absolute deviation of the firing latency across trials. Neurons that showed no/little response and whose jitter could not be determined were eliminated from further analysis. Neurons were annotated as showing somatic responses if they showed activation in the 0-6 ms window after light onset and had jitter of < 2 ms across trials when stimulated at 1 Hz with either 15 ms long square pulses or 20 ms long ramped pulses with 10 ms onset ramp in mice showing strong photoelectric voltage at light onset. Neurons that were activated in this window, but showed jitter > 2 ms were considered to be synaptically activated. Light responsivity was also assessed in later light ON (5-14 ms) or light OFF (16-25 ms) periods when the neurons were optogenetically stimulated at 1 Hz with 15 ms square pulses. In these periods light-responsive neurons were annotated as being synaptically activated or as synaptically inhibited in comparison to baselines. Neurons whose significance of response in the time window assessed could not be determined with the Mann-Whitney U test due to tied ranks were marked as unresponsive. To assess for the significance in the distribution of neurons showing a certain synaptic response across the behavioral groups (*Assessment+, Flight+* and *Others/Unresponsive*) a contingency table was created with the responsive neurons and the complement neuron numbers for the groups and tested using Fisher’s test, with Benjamini-Hochberg adjustment for pairwise comparisons between groups (*p_adj_ < 0.05*; Mangiafico, 2026).

To determine the decay time of the optogenetically-evoked response among the dPAG projecting VMHvl core neurons, z-scored mean PSTH’s across 1 Hz stimulation trials were created centered on the light onset and with bin width of 0.2 ms. The baseline of a mean PSTH was defined as the average response in the 45-60 ms period after light onset. Each mean PSTH was filtered (4th order Butterworth low-pass filter with 375 Hz cutoff; Van Boxtel, G.J.M., et al., 2021) and the time difference between the peak response and when the firing reached the PSTH baseline value after maximal response was calculated.

To investigate the intrinsic firing properties of the functional classes autocorrelogram shapes were assessed in the recording period when the behavioral test was carried out. 0-300 ms lags of counts autocorrelogram histogram with 1 ms bins were created and those *Assessment+* and *Flight+* neurons which showed a nonlinear decrease were selected to calculate the decay of intrinsic firing. An exponential decay function was fitted to filtered spike counts autocorrelogram (4th order Butterworth low-pass filter with 125 Hz cutoff; Van Boxtel, G.J.M., et al., 2021). Nonlinear least squares regression (Elzhov et al., 2023) was applied using equation 1: y(t) = A * exp(-t/τ) + C, or equation 2: y(t) = A * exp(-t/τ) * (1 + D/t * cos(ωx)) + C for neurons with prominent oscillations which caused a poor fitting with the first equation. The decay time constant, τ and autocorrelogram baseline C (normalized to spikes/s) were compared across the two functional classes using Mann-Whitney U test (*p < 0.05*). In case fitting with the exponential decay failed because the neurons did not show a nonlinear decrease, linear regression was used to find the slope of the autocorrelogram. The homogeneity of variances of τ between *Assessment+* and *Flight+* neurons was checked with the Fligner-Killeen test due to deviation of the distributions from normality (*p < 0.05*). To assess the distribution of the neurons showing a rapid exponential decay (τ ≤ 30 ms) in their autocorrelograms between the functional classes Chi square or Fisher test (if the count in any condition was < 5) for independence was carried out (*p < 0.05*). Neurons were assessed to be in the VMHvl core if they were found to be ChR2+ in the experiment where VMHvl neurons were opto-tagged by backlabelling from the dPAG, or ChR2- in the experiment where *Vgat+* neurons of the VMHvl were opto-tagged. Neurons were assessed to be in the VMHvl shell if they were found to be ChR2+ in the experiment where *Vgat+* neurons of the VMHvl were opto-tagged. While subclassifying neurons based on the optrode responses, those that were eliminated from the optogenetic stimulation analysis were also removed here.

To assess the regularity in firing of the neurons the coefficient of variation (CV) was determined using the formula CV = σ_ISI_/μ_ISI_. CV < 1 signifies neuronal firing that is more regular than a Poisson process while CV > 1 signifies less regular firing. Windows were created either 2 s after onset of flight or 2 s prior to the offset of investigation and used to calculate the behavior aligned CV for each unit. The measures were compared between the functional classes using the Mann-Whitney U test (*p < 0.05*).

Putative synaptic connections among the neurons of the behaviorally responsive clusters were determined with cross-correlograms. To assess whether observed cross-correlograms reflected significant temporal structure beyond chance, we compared each empirical cross-correlogram to a null distribution generated from the same unit pair. The shift predictor correlograms provided a null distribution against which to evaluate statistical significance and extract the bin-wise mean activity of the unit pair. For a given bin B in the empirical cross-correlogram, we denote the probability that the bin exceeds a chosen threshold *thr* under the null distribution by *P(B > thr)*. However, because each correlogram contains many bins, we tested a family-wise null hypothesis stating that no bin in the correlogram exceeds the threshold. For a cross-correlogram of *n* bins this joint null hypothesis can be written as:

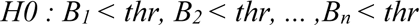

The condition *B_1_ < thr, B_2_ < thr, …,B_n_ < thr* is equivalent to the condition *B_max_ < thr* where *B_max_ = max_i_ {B_i_}* is the maximum bin value in the correlogram. This means:

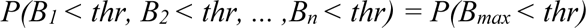

Thus, the distribution of *B_max_* under the null condition provides a direct way to control family-wise distribution. We estimated this distribution by computing the maximum bin value for each of the 750 random correlograms yielding a null distribution of maxima. We performed an analogous procedure for minima by computing *B_min_ = min_i_ {B_i_}* for each random correlogram allowing us to detect significant minima as well as maxima. For both *B_max_* and *B_min_*, we extracted the 99^th^ and 1^st^ percentiles of their respective null distributions. An empirical correlogram was considered significant if its maximum exceeded the 99^th^ percentile or its minimum fell below the 1^st^ percentile of the corresponding null distribution. Based on the symmetry of the response around histogram center and the lag of response in the shift-predictor subtracted histogram the unit pairs were manually annotated for the following connection types: fast (putatively monosynaptic) connections, slow connections, or receive common input, either excitatory or inhibitory in nature. The pairs with an asymmetric response centered around zero lag which crossed either side of the confidence interval (CI) were annotated to have a fast, putative monosynaptic connection when the lag was within 0-5 ms window and a slow connection when the lag was within the 5-25 ms window. When the response increased beyond the upper CI, it was considered to be an excitatory connection and when it decreased below the lower CI, an inhibitory connection. The response was considered to be led by the reference neuron when it occurred with positive lag and led by the target neuron when it occurred with negative lag, and were annotated independently. Therefore, each unit pair had two possible responses to these connection types. The pairs with responses symmetric around zero lag were considered to receive common input when the activity increased or decreased consistently across bins from the null distribution derived mean, without necessarily crossing either confidence interval. Each pair had a single possible response for the common input connections. To test the distribution of the connections among the like pairs (*Assessment+ - Assessment+*, A-A and *Flight+ - Flight+*, F-F) and unlike pairs (*Assessment+ - Flight+*, A-F and *Flight+ - Assessment+*, F-A) Chi square or Fisher test for independence was carried out for each connection type (*p < 0.05*).

### Histology

Serial coronal sections (40-50 µm) of perfused brains (1× PBS, 4% PFA in 1× PB) were cut on a cryo-microtome. For tracking final electrode placement brain slices were stained by immunostaining with DAPI (1:1000) or using a mounting medium with DAPI (MOWIOL with 5 mg/mL DAPI or Fluoromount). Images were taken with a fluorescence microscope (Leica or Slide Scanner) under 10-20× magnification to check the viral infection and verify optrode placement.

## Acknowledgements

We thank the EMBL Rome Laboratory Animal Facility (LAR), Gene Editing & Virus Facility (GEVF), Light Imaging Facility (LIF), and Roberto Voci and Valerio Rossi for support with mouse husbandry. This research was supported by European Molecular Biology Laboratory (EMBL) core funding, an EMBL PhD Fellowship to S.D., and a European Research Council (ERC) Advanced Grant (AdG) TERRITORY #101097411 to C.T.G.

## Author Contributions

S.D. and C.T.G. designed the research; S.D., E.W. and L.S. performed experiments with *in vivo* electrophysiology support from M.E.M.; S.D., S.T., E.W., S.K., S.A. and Y.Z. analysed the data with input from H.A.; S.D., S.T. and C.T.G. wrote the manuscript.

## Supplementary Figure Legends

**Figure S1.**
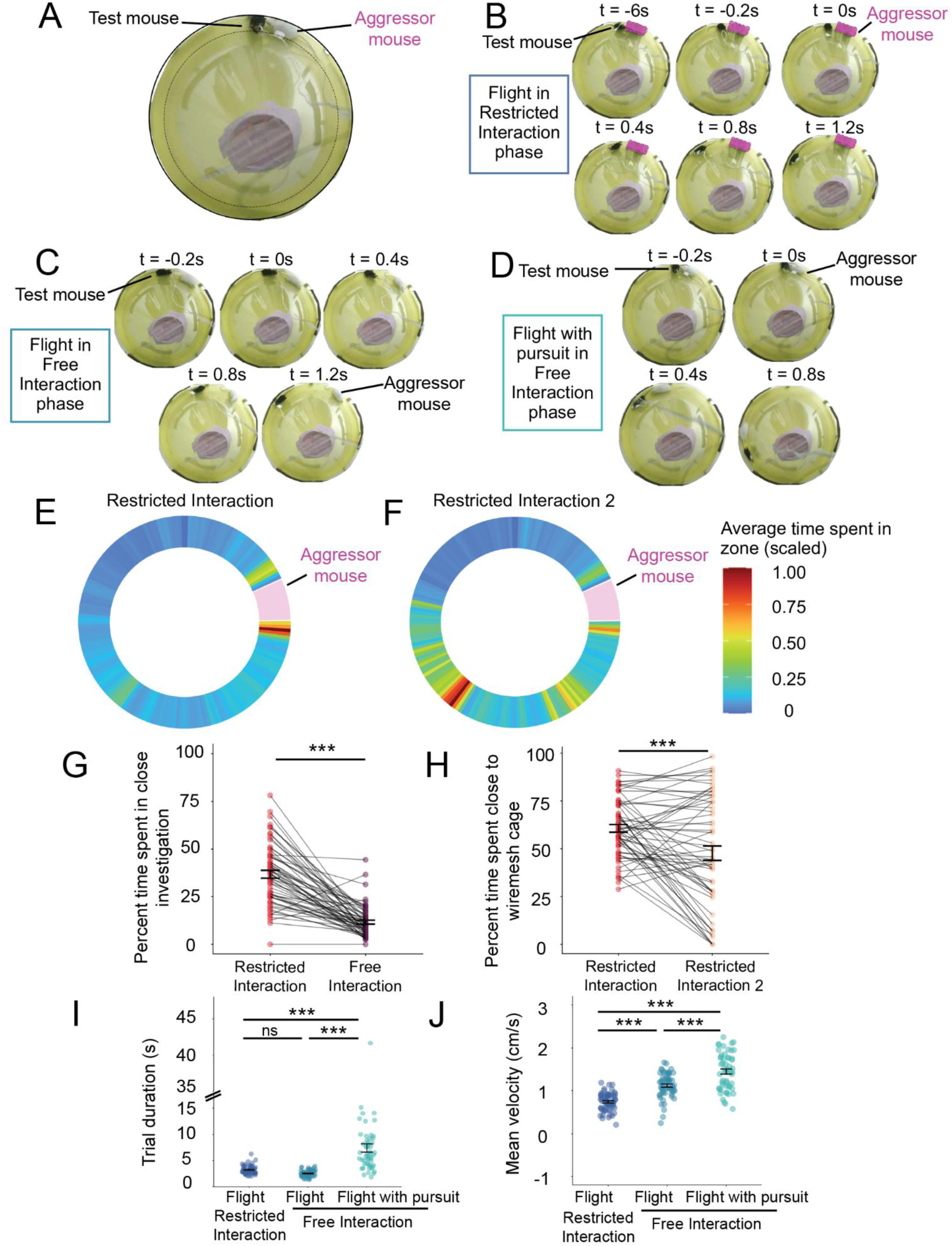
***A***, Top view of the circular arena during Free interaction phase. Social interaction occurs in the circular corridor bound by the outer wall (solid outer circle) and the inner cone (dashed circle). ***B***, Representative time lapse of a flight trial in the restricted interaction (RI) phase. In the pre-flight period the test mouse investigates the caged aggressor (magenta box) before transitioning to avoidance. Time from flight onset is indicated in seconds. ***C***, Representative time lapse of a flight trial in free interaction (FI) phase. ***D***, Representative time lapse of a flight with pursuit trial in free interaction (FI) phase. ***E***, Scaled heatmap of time spent in the circular corridor in the pre-defeat RI phase. Data averaged from a subset of behavioral sessions (N = 19 sessions, sector width = 2 deg). Sector of the circular corridor with the wiremesh cage with aggressor mouse coloured in magenta. ***F***, Scaled heatmap of time spent in the circular corridor in the post-defeat RI2 phase from the same sessions. ***G***, Comparison of percent time spent in close investigation in the RI vs FI phases. (Wilcoxon signed rank test, V = 1796, *p < 0.001*; error bars represent mean ± SEM). ***H***, Comparison of percent time spent in close investigation in the RI vs RI2 phases (Welch’s t-test, t = 3.74, df = 59, *p < 0.001*; error bars represent mean ± SEM). ***I***, Quantification of the trial duration of flights in RI (N = 59), flights in FI (N = 58) and flights with pursuit in FI (N = 54). Each point in a group is an average of the trial measures in a session (ANOVA F = 35.73, *p < 0.001*, Tukey’s flight_RI_ vs flight_FI_ *p = 0.5,* flight_RI_ vs flight with pursuit *p < 0.001*; flight_FI_ vs flight with pursuit *p < 0.001*; error bars represent mean ± SEM). ***J***, Quantification of the mean velocity of a trial for flights in RI (N = 59), flights in FI (N = 58) and flights with pursuit in FI (N = 54). Each point in a group is an average of the trial measures in a session (ANOVA F = 69.638, *p < 0.001*, Tukey’s flight_RI_ vs flight_FI_ *p < 0.001,* flight_RI_ vs flight with pursuit *p < 0.001*; flight_FI_ vs flight with pursuit *p < 0.001*; error bars represent mean ± SEM; * *p < 0.05, ** p < 0.01, *** p < 0.001*).

**Figure S2.**
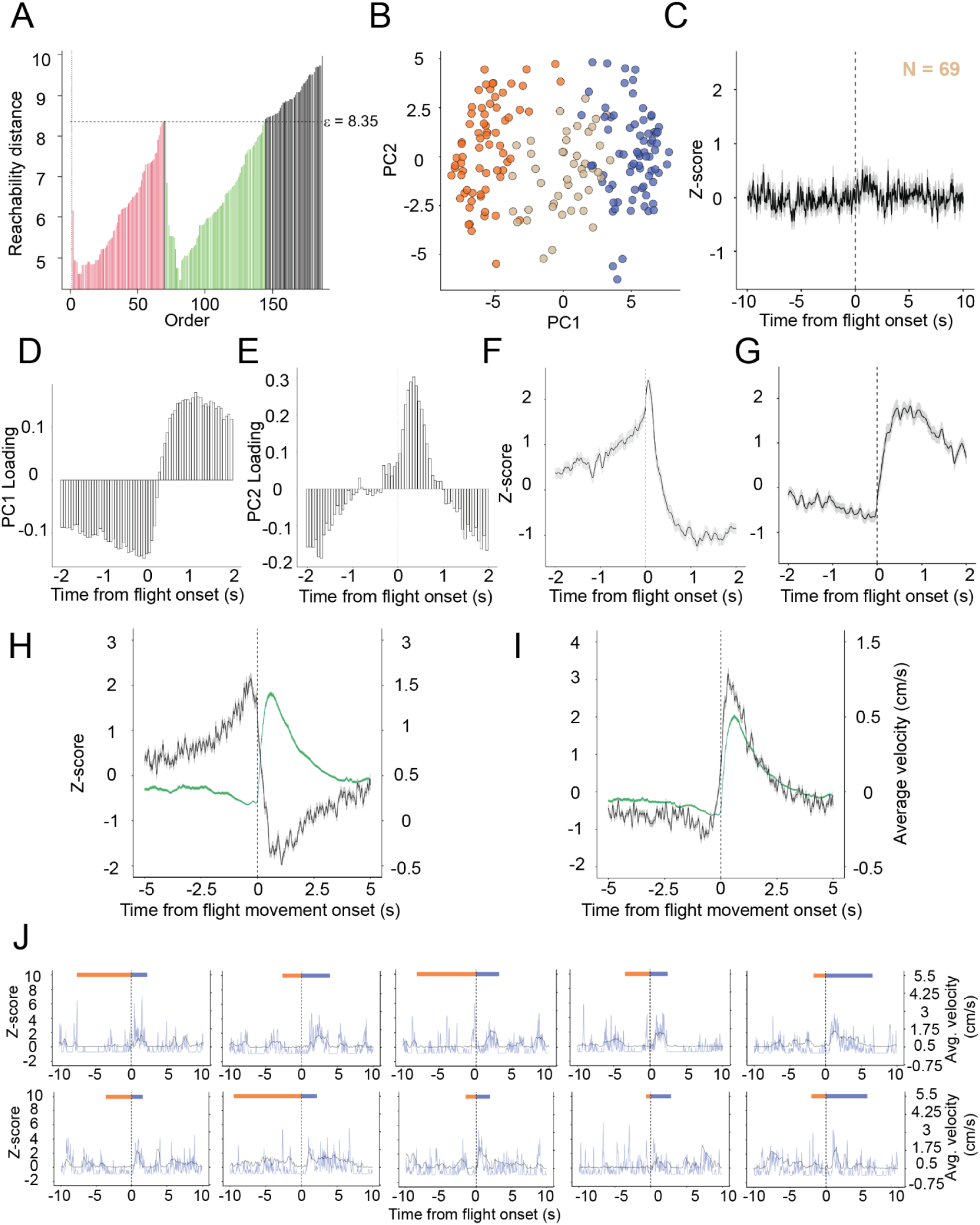
***A***, Clustering by OPTICS method using reachability plot. Plot of reachability distance against datapoints (behaviorally responsive neurons) ordered according to connectivity with neighbours. Cut-off of reachability distance e = 8.35 was applied to obtain cluster 1 (red) and cluster 2 (green) along with unclustered noise (black). ***B***, PCA projection of average PSTHs of the behaviorally responsive neurons coloured according to behavioral identity: *Assessment+* (orange)*, Flight+* (blue) and *Other* (tan). ***C***, Population average PSTH of the behaviorally responsive neurons not assigned to either functional class (*Others*, N = 44 neurons; grey shading indicates ±SEM). ***D***, Contribution of neural activity in the time bins of the PSTHs towards the PC1 dimension. ***E***, Contribution of neural activity in the time bins of the PSTHs towards the PC2 dimension. ***F***, 4 s population average PSTH of *Assessment+* neurons showing the peak of neural activity and transition after flight onset (grey shading indicates ±SEM). ***G***, 4 s population average PSTH of *Flight+* neurons (grey shading indicates ±SEM). ***H***, 10 s population average PSTH of *Assessment+* neurons centered on the onset of movement after manually annotated flight behavior start (black trace) along with the average velocity of flight (green trace; N = 69 neurons; shading indicates ±SEM). ***I***, 10 s population average PSTH of *Flight+* neurons centered on the onset of movement (black trace) with the average flight velocity (green trace; N = 74 neurons; shading indicates ±SEM). ***J***, Ten investigation-flight trials sampled from both Restricted Interaction (RI) and Free Interaction (FI) phases depicting 20s PSTH’s of the z-scored activity of a *Flight+* neuron (blue trace) and velocity (black trace) of the mouse, centered on the manually annotated flight behavior onset. The *Flight+* neuron is the same representative neuron as in **Figure 2D**.

**Figure S3.**
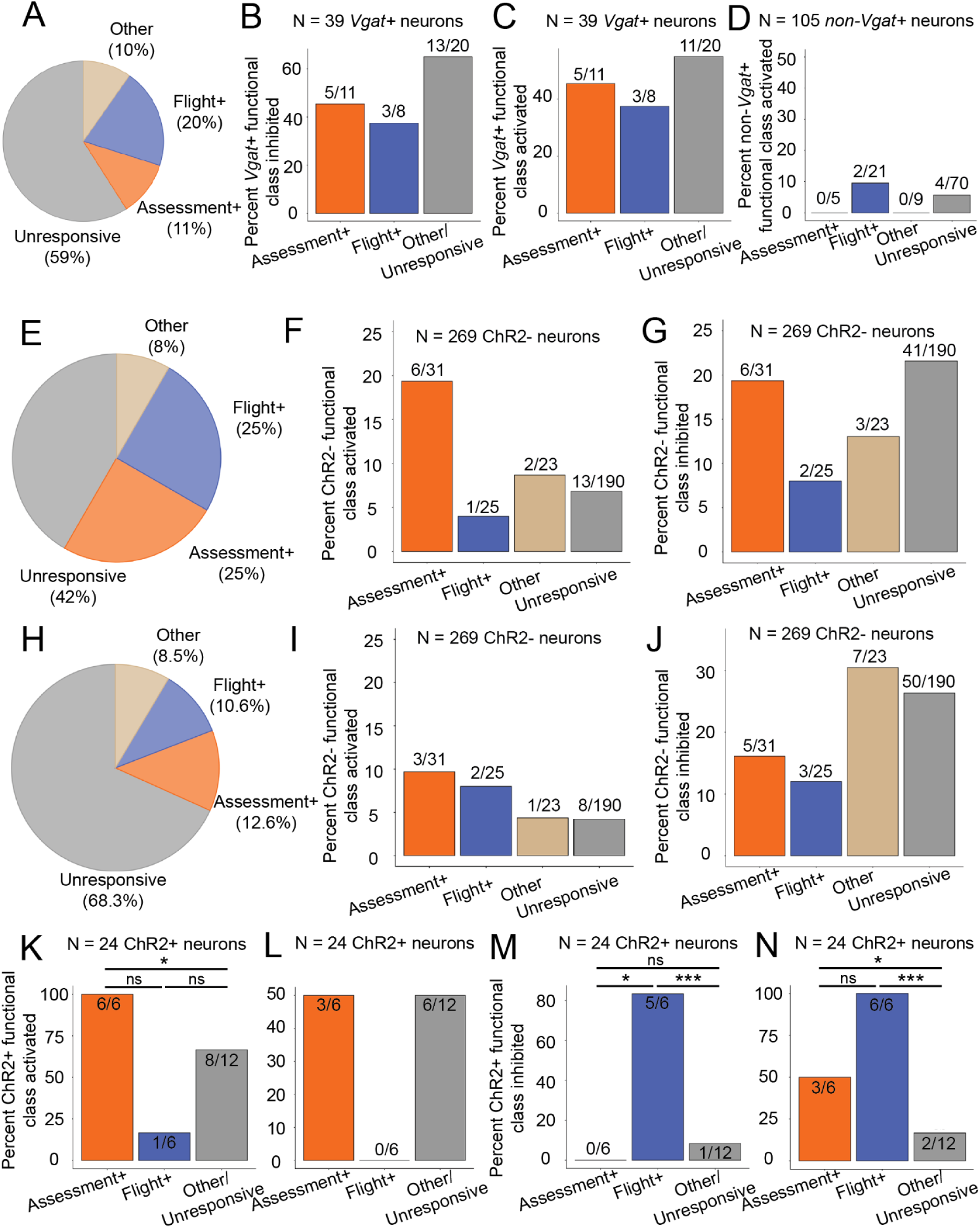
***A***, Distribution of functional classes recorded while opto-tagging *Vgat*+ neuron population in VMHvl (*Assessment+*: 16/144, *Flight+*: 29/144, *Others*: 14/144 and behaviorally unresponsive: 85/144). ***B***, Distribution of *Vgat*+ neurons receiving synaptic inhibition during the immediate post-light stimulation period (16-25 ms after light onset) among behavioral classes (N = 39 *Vgat*+ neurons; pairwise Fisher’s test: *Assessment+* vs *Flight+ p_adj_ = 1.00*, *Assessment+* vs *Others/Unresponsive p_adj_ = 0.674*, *Flight+* vs *Others/Unresponsive p_adj_ = 0.674*). ***C***, Distribution of *Vgat*+ neurons showing activation in the light stimulation period (5- 14 ms after light onset) among behavioral classes (N = 39 *Vgat*+ neurons; pairwise Fisher’s test: *Assessment+* vs *Flight+ p_adj_ = 1.00*, *Assessment+* vs *Others/Unresponsive p_adj_ = 1.00*, *Flight+* vs *Others/Unresponsive p_adj_ = 1.00*. ***D***, Distribution of non-*Vgat*+ neurons receiving synaptic activation during the immediate post-light stimulation period (16-25 ms after light onset) among behavioral classes (N = 105 non-*Vgat*+ neurons; pairwise Fisher’s test: *Assessment+* vs *Flight+ p_adj_ = 1.00*, *Assessment+* vs *Others/Unresponsive p_adj_ = 1.00*, *Flight+* vs *Others/Unresponsive p_adj_ = 1.00*). ***E***, Distribution of ChR2+ neurons among behavioral classes recorded while opto-tagging dPAG-projecting neurons in VMHvl (neurons with somatic response: N = 24, total neurons: N = 293; *Assessment+*: 6/24, *Flight+*: 6/24, *Others*: 2/24, *Behaviorally unresponsive*: 10/24; pairwise Fisher’s test: *Assessment+* vs *Flight+ p_adj_ = 0.76*, *Assessment+* vs *Others/Unresponsive p_adj_ = 0.04*, *Flight+* vs *Others/Unresponsive p_adj_ = 0.03*). ***F***, Distribution of ChR2- neurons receiving synaptic activation during the light stimulation period (5-14 ms after light onset) among behavioral classes (N = 269 neurons; pairwise Fisher’s test: *Assessment+* vs *Flight+ p_adj_ = 0.17*, *Assessment+* vs *Others/Unresponsive p_adj_ = 0.10*, Flight+ vs *Others/Unresponsive p_adj_ = 1.00*; pairwise Fisher’s test for all responsive neurons in the light stimulation period: *Assessment+* vs *Flight+ p = 0.01*)*. **G***, Distribution of ChR2- neurons receiving synaptic inhibition during the light stimulation period (5-14 ms after light onset) among behavioral classes (N = 269 neurons; pairwise Fisher’s test: *Assessment+* vs *Flight+ p_adj_ = 0.41*, *Assessment+* vs *Others/Unresponsive p_adj_ = 1.00*, *Flight+* vs *Others/Unresponsive p_adj_ = 0.41*). ***H***, Distribution of functional classes recorded while opto-tagging dPAG-projecting neurons in VMHvl (*Assessment+* = 37/293, *Flight+* = 31/293, *Others* = 25/293 and *Behaviorally unresponsive* = 200/293). ***I***, Distribution of ChR2- neurons receiving synaptic activation during the immediate post-light stimulation period (16-25 ms after light onset) among behavioral classes (N = 269 neurons; pairwise Fisher’s test: *Assessment+* vs *Flight+ p_adj_ = 1.00*, *Assessment+* vs *Others/Unresponsive p_adj_ = 0.48*, *Flight+* vs *Others/Unresponsive p_adj_ = 0.48*). ***J***, Distribution of the ChR2- neurons receiving synaptic inhibition during the immediate post-light stimulation period (16-25 ms after light onset) among behavioral classes (N = 269 neurons; pairwise Fisher’s test: *Assessment+* vs *Flight+ p_adj_ = 0.72*, *Assessment+* vs *Others/Unresponsive p_adj_ = 0.40*, *Flight+* vs *Others/Unresponsive p_adj_ = 0.40*). ***K***, Distribution of ChR2+ neurons activated during the light stimulation period (5-14 ms after light onset) among the behavioral classes (N = 24 ChR2+ neurons; pairwise Fisher’s test: *Assessment+* vs *Flight+ p_adj_ = 0.17*, *Assessment+* vs *Others/Unresponsive p_adj_ = 0.02*, Flight+ vs Others/Unresponsive *p_adj_ = 1.00*). ***L***, Distribution of ChR2+ neurons activated during the immediate post-light stimulation period (16-25 ms after light onset) among behavioral classes (pairwise Fisher’s test: *Assessment+* vs *Flight+ p_adj_ = 0.36*, *Assessment+* vs *Others/Unresponsive p_adj_ = 0.35*, *Flight+* vs *Others/Unresponsive p_adj_ = 1.00*). ***M***, Distribution of ChR2+ neurons inhibited during the light stimulation period (5-14 ms after light onset) among behavioral classes (pairwise Fisher’s test: *Assessment+* vs *Flight+ p_adj_ = 0.02*, *Assessment*+ vs *Others/Unresponsive p_adj_ = 1.00*, *Flight+* vs *Others/Unresponsive p_adj_ < 0.001*). ***N***, Distribution of ChR2+ neurons inhibited during the immediate post-light stimulation period (16-25 ms after light onset) among behavioral classes (pairwise Fisher’s test: *Assessment+* vs *Flight+ p_adj_ = 0.28*, *Assessment+* vs *Others/Unresponsive p_adj_ = 0.03*, *Flight+* vs *Others/Unresponsive p_adj_ < 0.001*; * *p < 0.05, ** p < 0.01, *** p < 0.001*).

**Figure S4.**
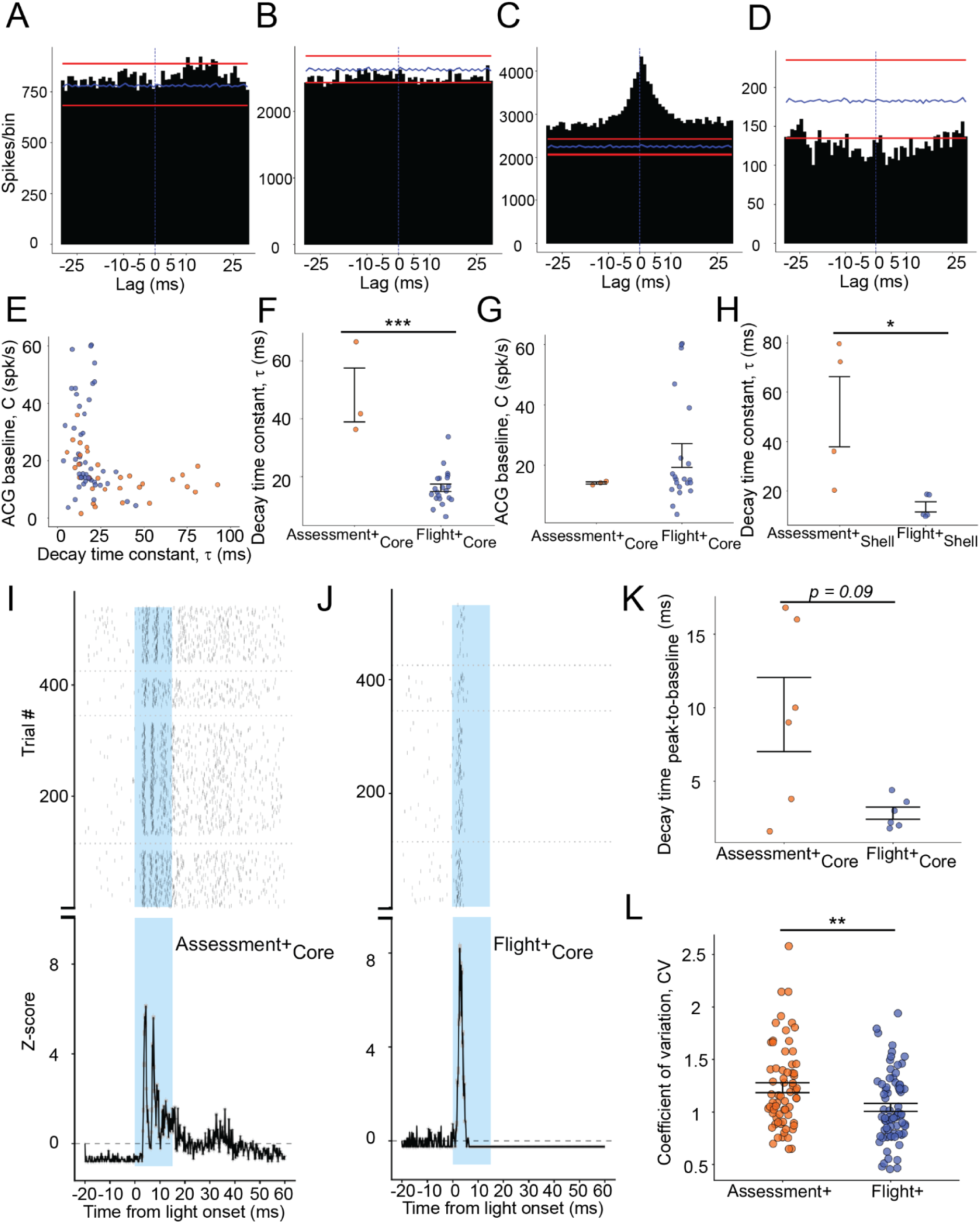
***A***, Representative example of a cross-correlogram between a pair of *Assessment+* neurons (A-A) exhibiting a slow (> 5 ms lag) excitatory connection (bin size = 1 ms). Blue line denotes the bin-wise mean count/bin and red lines indicate 1% and 99% confidence intervals derived from the null distribution. ***B***, Representative example of a cross-correlogram between an *Assessment+* and *Flight+* neuron showing a slow inhibitory connection. ***C***, Representative example of a cross-correlogram between a pair of *Assessment+* neurons receiving common excitatory input. ***D***, Representative example of a cross-correlogram between an *Assessment+* and *Flight+* neuron receiving common inhibitory input. ***E***, Distribution of the decay time constant τ against the autocorrelogram baseline C extracted from the exponential fitting of autocorrelograms of *Assessment+* (N = 32/69, orange) and *Flight+* (N = 45/74, blue) neurons (Spearman correlation – *Assessment+*: ρ = -0.56, *p < 0.001*; *Flight+*: ρ = -0.30, *p = 0.038*)*. **F**,* Comparison of decay time constant τ of functional classes in VMHvl core (*Assessment+,* N = 3 vs Flight+, N = 21; Mann-Whitney U test, W = 63, *p < 0.001*; Fligner-Killeen test χ^2^(1) = 1.00, *p = 0.317*; error bars represent mean ± SEM). ***G,*** Comparison of autocorrelogram baseline C of functional classes in VMHvl core (Mann-Whitney U test, W = 22, *p = 0.451*; error bars represent mean ± SEM). ***H,*** Comparison of decay time constant τ of functional classes in VMHvl shell (*Assessment+,* N = 4 vs Flight+, N = 5; Mann-Whitney U test, W = 20, *p = 0.015*; Fligner-Killeen test χ^2^(1) = 5.81, *p = 0.015*; error bars represent mean ± SEM). ***I,*** Representative example of the opto-response of an *Assessment+* unit expressing channelrhodopsin recorded while opto-tagging dPAG-projecting neurons in VMHvl. (**top**) Raster plot of the firing of the unit across light pulses of different stimulation frequencies separated by dashed lines (from bottom: 1, 2, 5 and 10 Hz, bin width = 0.2 ms). (**bottom**) Average response PSTH across all trials. ***J,*** Representative example of the opto-response of a *Flight+* neuron expressing channelrhodopsin. (**top**) Raster plot of the firing of the unit across light pulses of different stimulation frequencies separated by dashed lines (from bottom: 1, 2, 5 and 10 Hz, bin width = 0.2 ms). (**bottom**) Average response PSTH across all trials. ***K,*** Comparison of the time required for the optogenetically-evoked response to decay to baseline from the peak response between functional classes (*Assessment+,* N = 6 vs *Flight+*, N = 6; Mann-Whitney U test, W = 29, *p = 0.093*; error bars represent mean ± SEM). ***L***, Distribution of coefficient of variation of neural activity in 2 s windows when behaviorally-evoked activation occurs for each functional class (*Assessment+*: 2 s before investigation offset, *Flight+*: 2 s after flight onset; *Assessment+,* N = 69 vs Flight+, N = 74; Mann-Whitney U test, W = 3282, *p = 0.004*; error bars represent mean ± SEM; * *p < 0.05, ** p < 0.01, *** p < 0.001*).

**Table S1.**
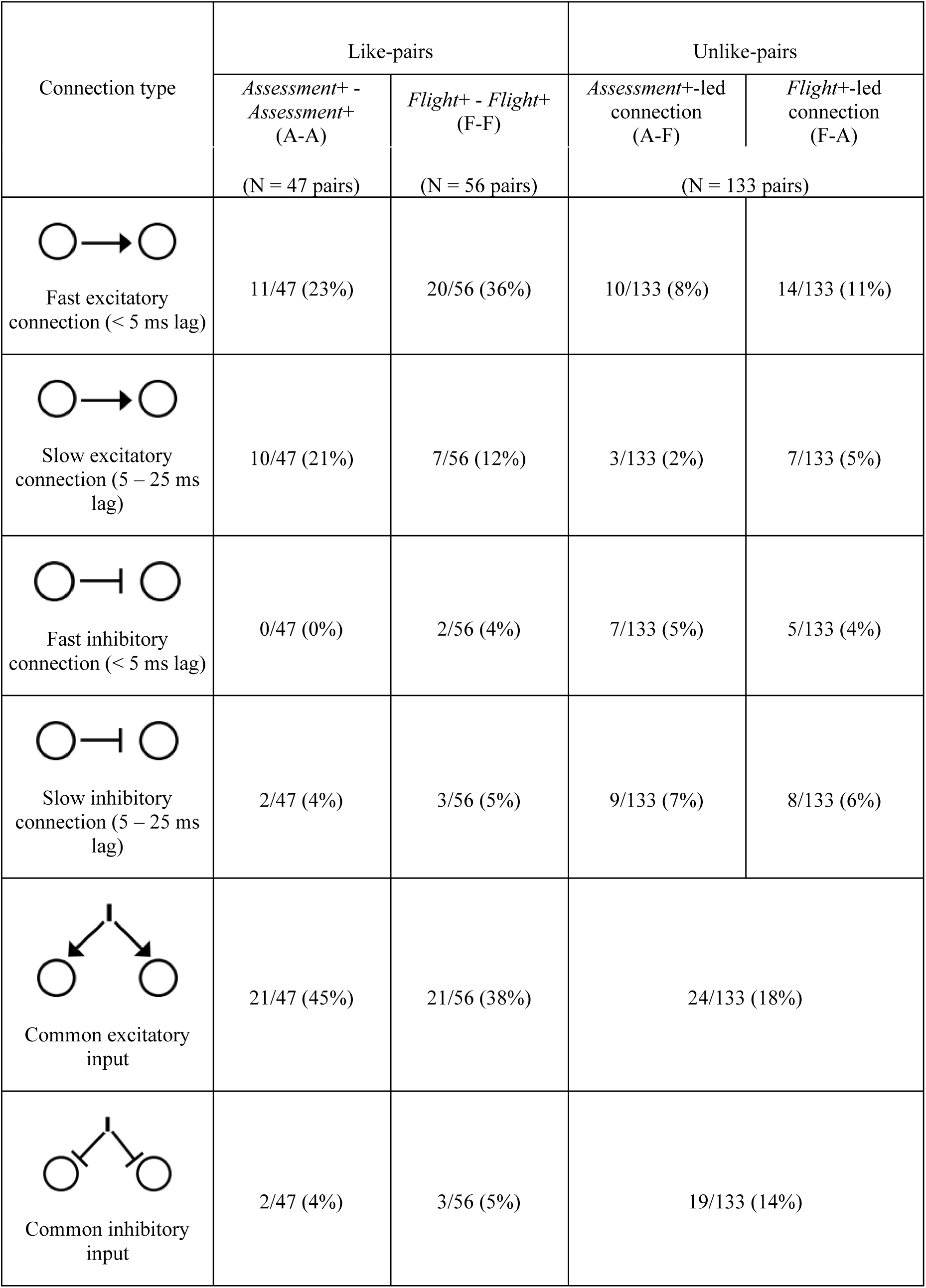
Table depicting the numbers of connections derived from cross-correlograms of like and unlike pairs of *Assessment+* and *Flight+* neurons in VMHvl.

| Connection type | Like-pairs |  | Unlike-pairs |  |
| --- | --- | --- | --- | --- |
|  | *Assessment+* - *Assessment+*<br>(A-A)<br>(N = 47 pairs) | *Flight+* - *Flight+*<br>(F-F)<br>(N = 56 pairs) | *Assessment+*-led<br>connection<br>(A-F)<br>(N = 133 pairs) | *Flight+*-led<br>connection<br>(F-A) |
| Fast excitatory<br>connection (< 5 ms lag) | 11/47 (23%) | 20/56 (36%) | 10/133 (8%) | 14/133 (11%) |
| Slow excitatory<br>connection (5 – 25 ms<br>lag) | 10/47 (21%) | 7/56 (12%) | 3/133 (2%) | 7/133 (5%) |
| Fast inhibitory<br>connection (< 5 ms lag) | 0/47 (0%) | 2/56 (4%) | 7/133 (5%) | 5/133 (4%) |
| Slow inhibitory<br>connection (5 – 25 ms<br>lag) | 2/47 (4%) | 3/56 (5%) | 9/133 (7%) | 8/133 (6%) |
| Common excitatory<br>input | 21/47 (45%) | 21/56 (38%) | 24/133 (18%) |  |
| Common inhibitory<br>input | 2/47 (4%) | 3/56 (5%) | 19/133 (14%) |  |

